# Progesterone receptor-A isoform interaction with RUNX transcription factors controls chromatin remodelling at promoters during ovulation

**DOI:** 10.1101/2021.06.17.448908

**Authors:** DT Dinh, J Breen, B Nicol, KM Smith, M Nicholls, A Emery, YY Wong, SC Barry, HHC Yao, RL Robker, DL Russell

## Abstract

Progesterone receptor (PGR) plays diverse roles in reproductive tissues and thus coordinates mammalian fertility. In the ovary, acutely induced PGR is the key determinant of ovulation through transcriptional control of a unique set of genes that culminates in follicle rupture. However, the molecular mechanisms for PGR’s specialised function in ovulation is poorly understood. To address this, we assembled a detailed genomic profile of PGR action through combined ATAC-seq, RNA-seq and ChIP-seq analysis in wildtype and isoform-specific PGR null mice. We demonstrated the unique action of PGR-A isoform in the ovary through a transcriptional complex involving physical interaction with RUNX and JUN/FOS transcription factors. The assembly of this unique complex directs targeting of PGR binding to proximal promoter regions and enables chromatin accessibility, leading to ovulatory gene induction. This PGR signalling mechanism is specific to ovulation and provides potential targets for infertility treatments as well as new contraceptives that block ovulation.

## 1. INTRODUCTION

Ovulation is a highly controlled process that ensures the release of appropriately mature oocytes, timed to coincide with the receptivity of the uterus for embryo implantation. The mechanism of ovulation begins with the activation of cAMP-dependent kinase (PKA) and extracellular regulated kinase (ERK) signalling cascades in granulosa cells of the follicle in response to the pituitary LH surge ^1^. These signals result in the induction of an array of transcriptional regulators that collectively mediate complete structural remodelling of the ovarian follicle and the resumption of oocyte meiotic maturation ^2^. Meanwhile, follicular granulosa cells undergo pervasive differentiation and rapid neoangiogenesis to form the corpus luteum, a transient endocrine organ that produces high levels of progesterone to promote uterine receptivity ^3^ and mammary gland maturation ^4^ in preparation for pregnancy and lactation.

Progesterone signalling also has an obligatory role in granulosa cells where it is essential for ovulation. In the ovary, PGR is a key transcription factor that determines ovulation ^5,6^, as demonstrated in mutant animal models as well as in primates and human. Female mice with either total or granulosa-specific null mutation of the nuclear progesterone receptor (PGR) have a complete anovulatory phenotype without affecting oocyte meiosis or luteinisation ^5–7^. Likewise, in primates and humans, administration of PGR antagonists (Mifepristone or Ulipristal Acetate) blocks ovulation ^8,9^ and thus have been explored as potential acute contraceptive agents. However, disrupting systemic progesterone action has unacceptable side effects across many target organs, necessitating further research into the unique ovarian PGR action and its downstream target genes in order to discover highly selective targets for ovulation-blocking contraceptives.

As a steroid receptor, PGR is a nuclear transcription factor that is activated by its ligand, progesterone. In the uterus and mammary gland, PGR expression is maintained by estrogen action. However, in the ovary PGR is not estrogen dependent ^10^ but is highly induced in granulosa cells in response to the LH surge ^11^. In general, PGR is believed to regulate transcription through binding the PGR response element (PRE) motif that can be recognised by all NR3C steroid receptors. However, PGR function can be highly intricate and involves binding to target genes in cooperation with other transcription factors ^12,13^. Such transcriptional complexes can recruit histone and chromatin remodellers including steroid receptor coactivators (SRC), CBP/p300 and PRMT1 ^14–16^ to facilitate accessibility for the basal transcriptional machinery and RNA Pol II to induce expression of target genes ^17^. The two major isoforms of PGR – PGR-A and PGR-B – possess specific properties and functions, including distinct protein-protein interaction capabilities ^18^ and cross-isoform regulation, as demonstrated by the widely reported repression of PGR-B transcriptional activity by PGR-A in the uterus ^19^ and breast cancer cells ^20^. In the ovary, both PGR isoforms are induced by the LH surge ^21,22^; however, PGR-A is credited as the more essential isoform in ovulation, as shown through PGR-A specific KO mouse model (AKO) ^23^. In contrast, female mice lacking only PGR-B (BKO) exhibit normal ovulation ^24^. The relative importance of PGR-A and PGR-B throughout reproductive physiology remains an active topic of investigation ^3,25,26^, and their molecular mechanisms in granulosa cells that control ovulation remain unknown.

Our previous characterisation of the granulosa cell PGR cistrome through ChIP-seq suggests that the classic pathway involving the canonical PRE motif is not favoured by PGR in granulosa cells and that the cooperation between PGR and context-specific transcription factors, such as JUN/FOS and RUNX transcription factor families, establishes unique transcriptional regulation during ovulation ^22^. The interaction between transcription modulators and PGR has been shown to be important in other biological contexts ^12,13^; however, whether such interactions are crucial for PGR-mediated ovarian functions remains unexplored. Among the potential binding partners for PGR, the RUNX family was uniquely identified in granulosa cells ^22^. Included in this family is RUNX1, which acts as a regulator of gene expression in peri-ovulatory granulosa cells ^27,28^. The ablation of CBFβ, the canonical protein partner of RUNX1 and the closely related RUNX2, results in the reduction of important ovulatory genes and consequentially impaired fertility in female mice ^29,30^. However, the nature of PGR/RUNX1 relationships in the context of ovulation remains unknown.

This study provides novel insights into the molecular mechanisms employed by PGR in granulosa cells to cause ovulation. The scope of global changes in the open chromatin landscape and transcriptome induced by the LH surge and PGR in mouse granulosa cells was defined using ATAC-seq and RNA-seq in wildtype and PGR-null mice. The formation of molecular complexes between PGR and other transcription factors was confirmed. For RUNX1, comparative ChIP-seq analysis pointed to a high level of PGR/RUNX1 cooperativity at mutual chromatin sites and especially at proximal promoters of ovulatory genes. Isoform-specific granulosa cell transcriptomes also identified the dominant role of PGR-A in ovulation, through interaction with RUNX transcription factors. The results demonstrate a unique molecular mechanism that involves PGR-A and RUNX1 co-operativity to shape granulosa cell chromatin epigenetic reprogramming and transcriptional response to the LH surge. This novel progesterone signalling pathway is critical for normal fertility and will enable the identification of causes of unexplained infertility as well as the discovery of molecular complexes or protein targets for future contraceptive discovery.

## 2. RESULTS

### Rapid global reprogramming of chromatin state and transcription by LH-stimulus through the induction and repression of distinct transcription factor binding

Ovulation induction by hCG injection to simulate the LH surge in mice provides an *in vivo* model for the rapid response to ovulatory cues in granulosa cells. ATAC-seq in granulosa cells isolated before or 6 h after *in vivo* LH-stimulus revealed dynamic changes in chromatin accessibility in response to the induction of ovulatory transcription factors, including PGR. Stringent ATAC-seq peak calling criteria identified a total of 71,287 peaks across both conditions, of which 28,729 sites (40.3%) were LH-induced, 21,257 sites (29.8%) were significantly repressed and 21,301 sites (29.8%) with no change in accessibility after LH-stimulus (Figure 1A). LH-mediated differentially accessible sites, including those more accessible before or after LH-stimulus, indicated that chromatin remodelling occurred not only in promoters but also at distal enhancers and throughout gene bodies (Figure 1B). LH-stimulus induced the accessibility of promoters corresponding to genes involved in cytoskeletal reorganisation, mitochondrial functions, and signalling pathway activation which are important processes leading up to ovulation (Supplementary figure 1). Significantly, LH-stimulus also repressed access to genes that were responsible for gene silencing and nucleosome modification. Genes that were involved in normal cellular processes, such as RNA processing, biosynthesis and vesicle transportation, showed no change in promoter accessibility by the treatment.

**Figure 1:**
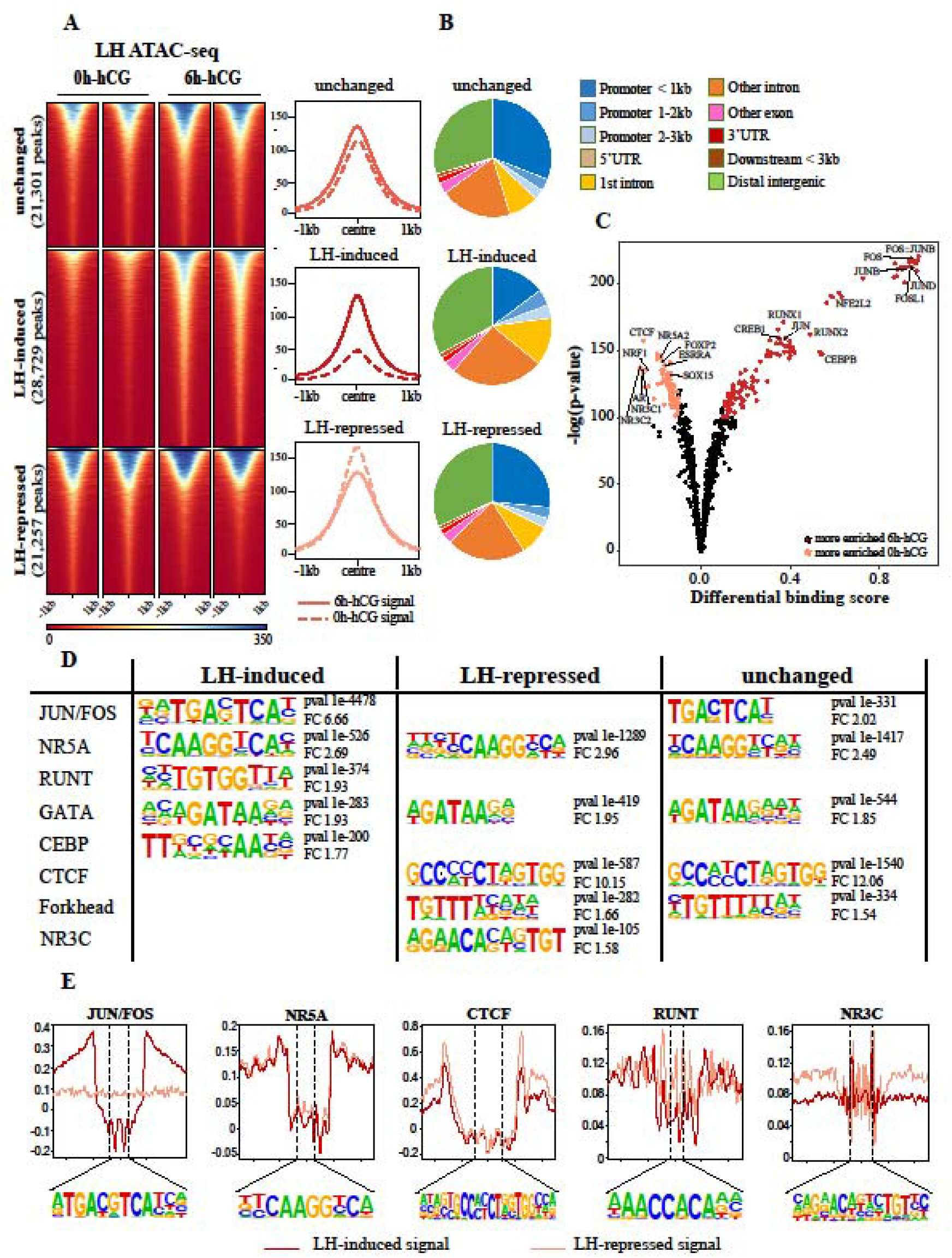
Global changes in the chromatin landscape of granulosa cells after hCG treatment. (A) Heatmap of LH ATAC-peak signals (left), with peaks subset into those that were unchanged (top), LH-induced (middle) or LH-repressed (bottom). Representative profile plots of each condition are on the right. (B) Genomic distribution of ATAC-peaks that were unchanged by LH, LH-induced or LH-repressed. (C) Differential enrichment analysis of transcription factor-binding motifs in ATAC-seq peaks found in 6h-hCG and 0h-hCG. Top 5% motifs that were differentially enriched in either condition are highlighted. (D) *de novo* motif enrichment analysis of LH-induced, LH-repressed or unchanged ATAC peaks as identified through HOMER. For each dataset, top 5 motifs are shown with p-value and fold change to background (FC). (E) DNA footprints of selected motifs in ATAC-peaks that were LH-induced (red) or LH-repressed (pink).

As ATAC-seq provides DNA accessibility data at footprint resolution, it can be used to analyse transcription factor binding sites in an unbiased manner. To predict the activity of key transcription factors involved in changes in the accessibility of LH-mediated open chromatin sites, differential enrichment analysis of transcription factor-binding motifs was performed (Figure 1C, Supplementary table 1). The motif for many ovulatory transcription factors, such as for JUN/FOS, RUNX, CREB and CEBPβ, showed significantly increased binding score in peaks for open chromatin after LH-stimulus, whereas those for CTCF, NR5A and ESRR were enriched at pre-LH open chromatin sites. Interestingly, the canonical motifs for the NR3C family members, such as glucocorticoid receptor (NR3C1/GR), mineralocorticoid receptor (NR3C2/MR), androgen receptor (NR3C4/AR) as well as PGR, were significantly more enriched in open chromatin before LH-stimulus. In support of these findings, *de novo* motif enrichment analysis using HOMER also identified JUN/FOS, RUNT and CEBP motifs in LH-induced, but not LH-repressed open chromatin regions (Figure 1D). Motifs for NR5A and GATA families were similarly enriched in all ATAC-seq peak subsets. Importantly, a motif best matched to that bound by NR3C members was identified only in the LH-repressed subset, consistent with the differential transcription factor binding analysis pattern above. Direct binding of transcription factors to these motifs was confirmed through DNA motif footprinting (Figure 1E). Footprint patterns indicative of transcription factors bound to these sites could be observed with medium to deep indentation (JUN/FOS, CTCF, NR5A) and shallower patterns (RUNT, NR3C), alluding to differences in the dwell time of these transcription factors. Again JUN/FOS footprints were only evident after LH-stimulus, while evidence of NR3C footprints was more obvious in unstimulated cells.

In order to correlate chromatin accessibility profiles with gene expression changes, the complete transcriptional response of peri-ovulatory granulosa cells to LH-stimulus was profiled through RNA-seq of granulosa cells collected before or 8 h after *in vivo* LH-stimulus. This timepoint was chosen as it is approximately 2 h after the peak induction of key ovulatory transcription factors. A total of 2,088 genes were differentially expressed (DEG), of which 52% were upregulated by LH-stimulus (Figure 2A). This included many genes previously associated with ovulation, such as *Ptgs2, Pgr, Runx1, Runx2* and *Adamts1* (Supplementary Table 2). Prediction of the potential upstream regulators of LH DEGs using tools from the Ingenuity Pathway Analysis (IPA) software identified a suite of transcription factors, many of which were also among the DEGs in response to the LH surge, including PGR (logFC = 6.4), RUNX1 (logFC = 4.6), CEBPβ (logFC = 2.5) and JUNB (logFC = 2.1) (Supplementary Table 2, 3). Notably, the binding motifs for these same transcription factors also showed increased accessibility following LH-stimulus (Figure 1C, D). The global relationship between promoter accessibility and downstream gene expression in peri-ovulatory granulosa cells was illustrated through a comparison between LH DEG and LH-dependent chromatin accessibility at the promoter (within 3 kb upstream of TSS) of these genes. Significant changes in promoter accessibility corresponded with transcriptional change in 69.3% of LH-driven DEGs (1,447 out of 2,088 genes) (Figure 2B, red symbols), in which promoter accessibility largely increased in LH-induced genes and was less accessible in LH-repressed genes (Spearman correlation coefficient = 0.467). For example, in *Cxcr4* and *Inhbb*, two genes that were respectively upregulated and downregulated post-LH, clear association between expression pattern and ATAC-seq signal at the gene promoter could be observed (Figure 2C). In some instances, however, genes that were differentially regulated had constitutively accessible promoters, as seen in *Cited1*.

**Figure 2:**
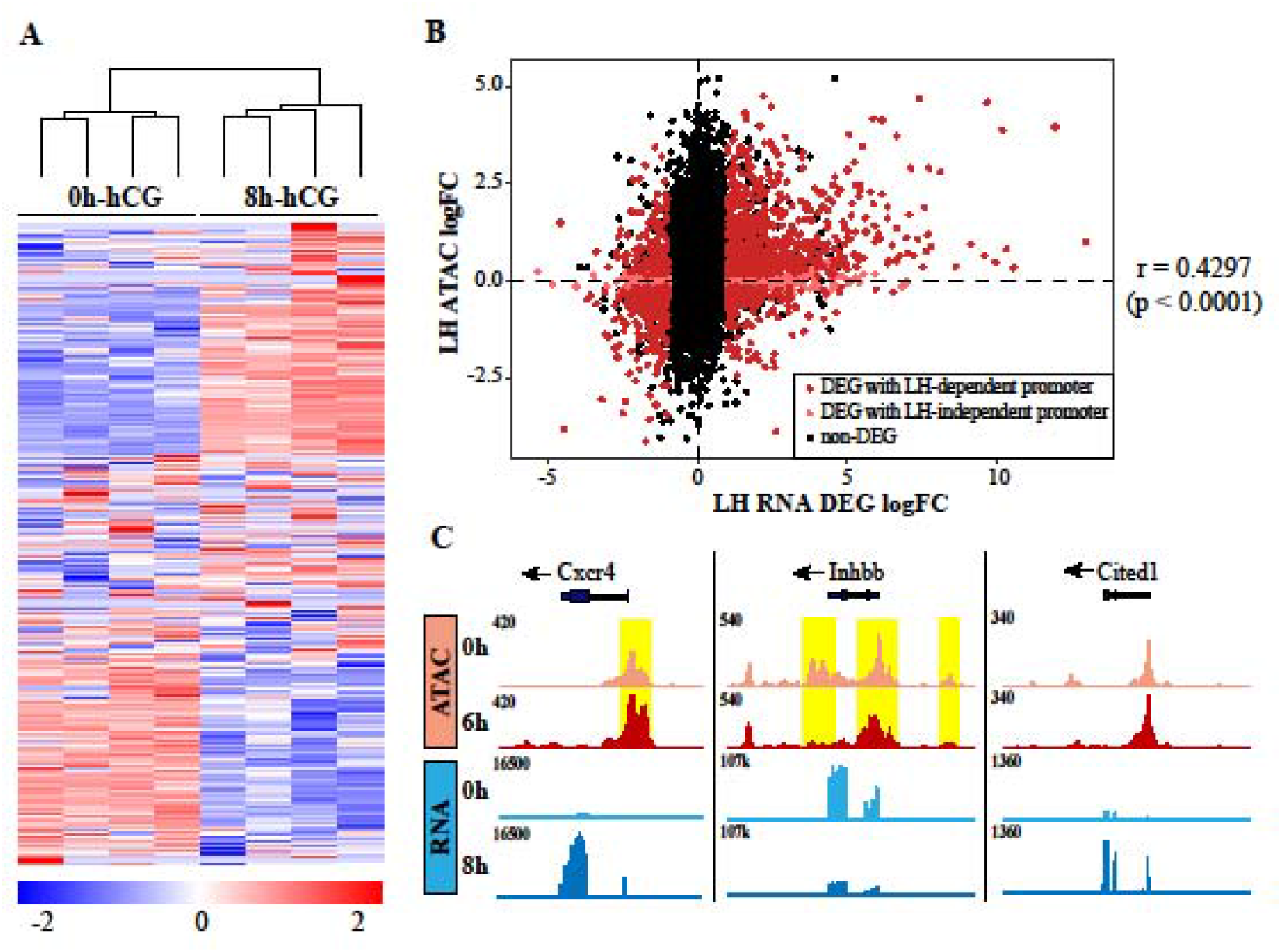
Global changes in the transcriptome of granulosa cells after hCG treatment. (A) Heatmap of the expression level of ovulatory DEG identified in LH RNA-seq. (B) Correlation between expression level and promoter accessibility at RNA-seq identified ovulatory genes, shown as LH DEG RNA-seq log fold change vs ATAC-seq log fold change at their respective promoter (within 3 kb of TSS). DEGs with LH-dependent promoter are labelled red, DEGs with LH-independent promoter are labelled pink. Correlation between RNA-seq and ATAC-seq intensity is determined through Spearman’s correlation coefficient (r = 0.4297, p < 0.0001) (C) Examples of ATAC-seq (track 1, 2) and RNA-seq (track 3, 4) signals for 0h-hCG (light) and 6h/8h-hCG (dark) at *Cxcr4, Inhbb* and *Cited1*. Differentially accessible ATAC peaks are highlighted in yellow.

### PGR and RUNX1 mutually bind chromatin in peri-ovulatory granulosa cells

Since RUNX1 is functionally important in granulosa cells ^31^ and RUNT motifs were enriched in granulosa cell PGR ChIP-seq peaks ^22^ as well as regions which gained chromatin accessibility after LH-stimulus (Figure 1C), we asked whether RUNX1 is key to the transcriptional regulatory mechanism that mediates ovulatory gene expression. Having confirmed RUNX1 expression to be induced in granulosa cells in response to the LH surge (Supplementary figure 2A, B), we sought to biochemically determine its targets on a genome-wide scale. RUNX1 ChIP-seq was performed on granulosa cells before or 6 h after *in vivo* hCG treatment to induce ovulation. Global RUNX1-chromatin binding was detected prior to the LH surge but was also induced 7-fold after the LH surge (16,065 LH-induced peaks) (Figure 3A). Such LH-activated RUNX1 binding was strongly associated with LH-induced open chromatin as defined through ATAC-seq (Supplementary figure 3A). A clear relationship between RUNX1 and PGR chromatin interaction in peri-ovulatory granulosa cells was demonstrated through comparison between RUNX1 ChIP-seq and PGR ChIP-seq at 6h post-LH stimulation. PGR and RUNX1 shared a remarkable number of mutual binding sites within 100 bp, with 9,704 chromatin sites (or 70% of total PGR and 52% of total RUNX1 binding sites) having both PGR and RUNX1 binding (Figure 3B). Among these shared PGR/RUNX1 binding sites, 9,288 were located in ATAC-seq open chromatin sites identified at the same time point. Dividing transcriptionally active RUNX1 and PGR cistromes into those that were shared or uniquely bound by each transcription factor showed a relatively low level of promoter binding in uniquely PGR-bound sites, yet a strong preference for proximal promoter binding (within 1 kb of TSS) was a predominant characteristic of PGR/RUNX1 co-binding sites (Figure 3C). The remarkably high overlap of PGR and RUNX1 binding sites, coupled with our previous report of enriched RUNX1-binding motifs in granulosa cell PGR ChIP-seq ^22^ suggests that the previously reported granulosa-specific ability of PGR to target proximal promoter relies on co-binding with RUNX1. Such dependency was not reciprocal, as RUNX1 showed a prevalence for proximal promoter occupancy that was independent of PGR co-binding. To test whether LH-induced RUNX1 chromatin binding was dependent on PGR, we assessed whether LH-induced RUNX1 binding intensity was greater at PGR-overlapped sites versus PGR-independent sites. No difference was observed in the level of RUNX1 enrichment, suggesting that PGR was unlikely to play a role in facilitating the tethering and recruitment of RUNX1 to these shared sites (Supplementary figure 3B).

**Figure 3:**
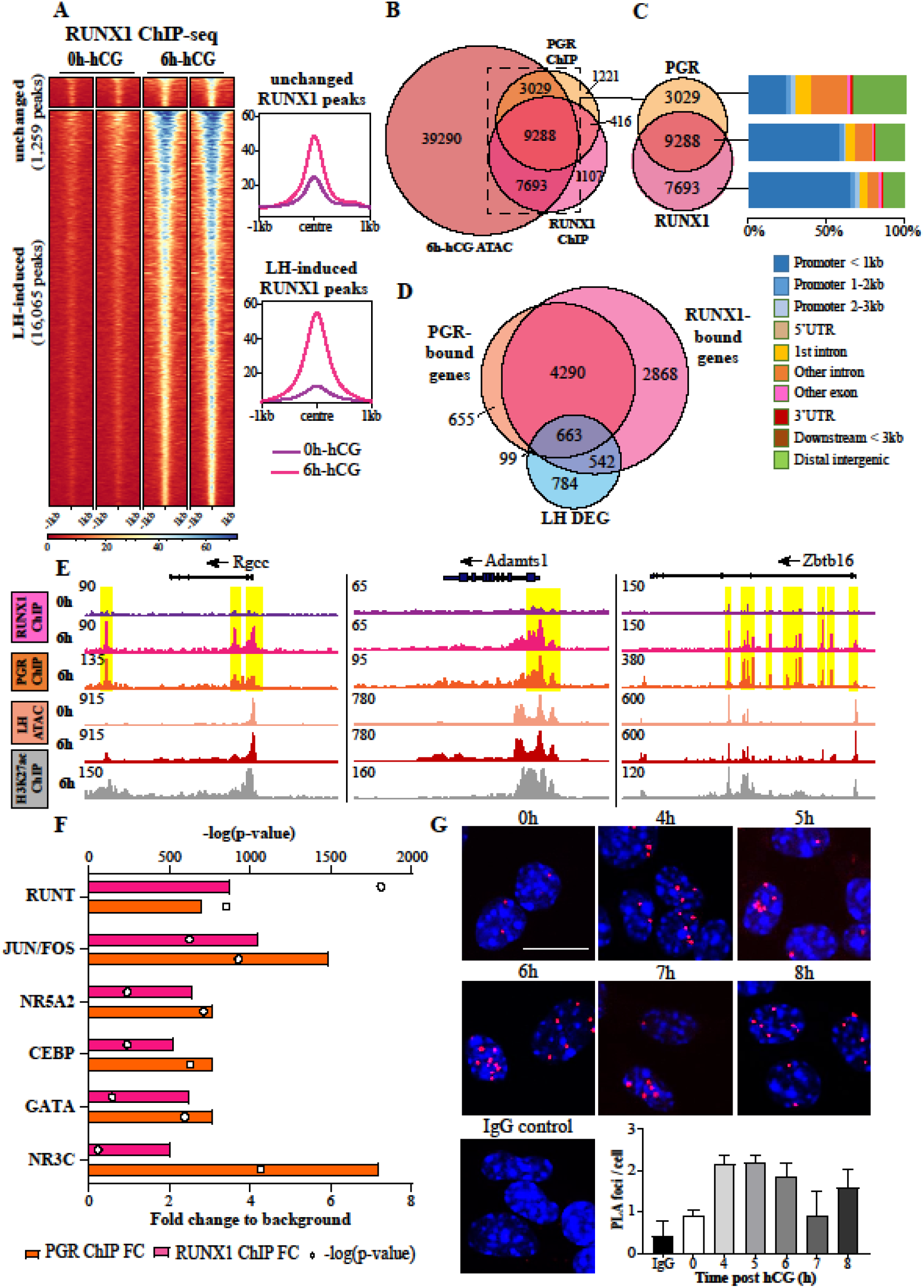
PGR and RUNX1 share mutual chromatin targets in peri-ovulatory granulosa cells. (A) Heatmap of RUNX1 ChIP-seq peak signals, with peaks subset into those that were unchanged (top) or LH-induced (bottom). Only 1 RUNX1 peak is LH-repressed and is thus not plotted (B) Venn diagram showing shared and factor-unique peak counts for RUNX1 and PGR in relation to 6h-hCG LH ATAC. (C) Genomic distribution of transcriptionally active PGR-specific (top), PGR/RUNX1 shared (middle) and RUNX1-specific (bottom) binding sites. (D) Venn diagram showing the relationship between LH DEG and genes with RUNX1 and/or PGR binding in their promoters. (E) Example of LH-dependent RUNX1 binding (track 1, 2) in conjunction with PGR (track 3) at *Rgcc, Adamts1* and *Zbtb16*. ATAC-seq peaks at 0h, 6h and H3K27ac ChIP-seq peaks at 6h (track 4-6) indicate open chromatin regions. (F) Top most common known sequence motifs found to be enriched at RUNX1 (pink) and PGR (orange) binding sites. Bars indicate fold enrichment of motif to background (bottom x axis). Circles indicate −log(p-value) (top x axis). (G) PLA for PGR/RUNX1 in cultured granulosa cells under hCG/R5020 time course treatment. One representative image is shown for each treatment. PGR/IgG antibody pair was used as negative control. PLA was performed in 3 biological replicates (4 mice per replicate). Red indicates protein-protein interaction. Blue is DAPI nuclear stain. Scale bar = 10 μm. Quantification of PLA signal is displayed as the number of nuclear foci per cell, displayed as mean ± SEM, p-value = 0.1044.

The importance of PGR and RUNX1 cooperation on ovulatory gene regulation at a genome-wide level was demonstrated by comparing genes with PGR and/or RUNX1 binding at their promoters against all DEG responding to LH in granulosa cell (Figure 3D). Remarkably, 62.4% of all LH-mediated DEGs contained at least PGR or RUNX1 binding. Of these, the majority of RUNX1- (55%) and PGR-bound (87%) LH DEGs were in fact co-bound by both transcription factors, implying that simultaneous interaction of both PGR and RUNX1 at target promoters was a major mechanism for each of these transcription factors in gene regulation. Examples of PGR/RUNX1 mutual chromatin occupancy were evident in the promoters of many known PGR and RUNX1 target genes in peri-ovulatory granulosa cells (*Adamts1, Rgcc* and *Zbtb16*) (Figure 3E).

DNA motifs preferentially bound by RUNX1 in granulosa cells were identified using motif enrichment analysis. As expected, the canonical RUNT motif was significantly highly enriched in the RUNX1 cistrome (3.5-fold over background, p-value 1e-787) as well as PGR cistrome (2.8-fold, p-value 1e-370) (Figure 3F). The NR3C motif was also enriched in RUNX1-bound peaks (2-fold over background, p-value 1e-25); interestingly, such enrichment was restricted to only PGR/RUNX1 co-binding sites (Supplementary figure 3C). Other enriched non-canonical motifs included those corresponding to JUN/FOS, CEBP and GATA transcription factors.

The potential physical interaction between PGR and RUNX1 was supported by positive proximity ligation assay (PLA) signals between PGR/RUNX1 in the nucleus of mouse granulosa cells treated with hCG and R5020 for up to 8 hours to mimic the *in vivo* ovulatory stimuli (Figure 3G). PLA signal showed a rapid transient induction of PGR/RUNX1 complexes by LH-stimulus, indicating an acute temporal PGR/RUNX1 interaction that was largely absent before stimulus but increased within 4 h after stimulation.

In addition to RUNX1, other members of the RUNT transcription factor family are capable of binding the RUNT motif. Notably, RUNX2 was induced by the LH surge in granulosa cells of antral follicles and had a transient temporal expression pattern during the peri-ovulatory window similar to PGR and RUNX1 (Supplementary figure 2A, B), consistent with previous reports ^32^. The interaction between PGR and RUNX2 was also demonstrated through PLA of granulosa cells exposed to ovulatory stimuli (Figure 4A). Similar to the interaction between PGR and RUNX1, PGR/RUNX2 PLA signal was induced within 4 h of stimulus. Additional potential candidate protein partners from the NR5A and JUN/FOS transcription factor families were selected based on their known involvement in granulosa cell functions during ovulation. Interactions of PGR with LRH1 and JUND were observed in granulosa cells after 6 h hCG, but not with cJUN or JUNB (Figure 4B). The interaction between PGR and ovulatory transcription factors was predominantly in the nuclear compartment of the cell, indicating that these transcription factors concurrently interact in chromatin-bound complexes in response to ovulatory cues.

**Figure 4:**
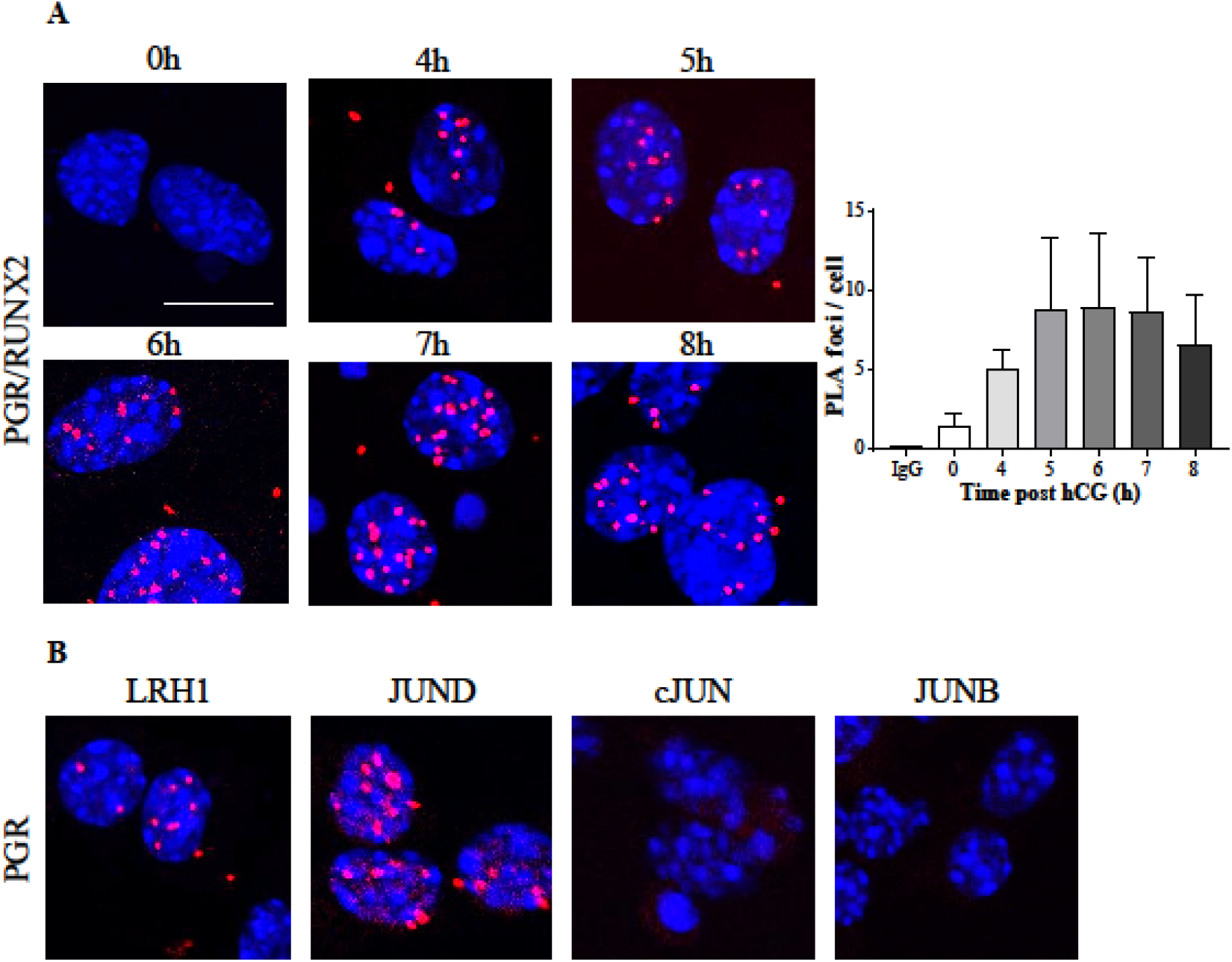
PGR interacts with other ovulatory transcription factors in granulosa cells in response to the LH surge. (A) PLA for PGR/RUNX2 in cultured granulosa cells under hCG/R5020 time course treatment. One representative image is shown for each treatment. PGR/IgG antibody pair was used as negative control. PLA was performed in 3 biological replicates (4 mice per replicate). Quantification of PLA signal is displayed as the number of nuclear foci per cell, displayed as mean ± SEM, p-value = 0.2294. (B) PLA showing the interaction between PGR and LRH1, JUND, cJUN and JUNB. Red indicates protein-protein interaction. Blue is DAPI nuclear stain. Scale bar = 10 μm.

### PGR mediates chromatin accessibility in granulosa cells

PGR-bound regions predominantly overlap with transcriptionally active chromatin, as shown by 75% overlap in PGR- and H3K27ac ChIP-seq peaks ^22^; likewise 90% of PGR binding sites were located in 6 h hCG ATAC-seq peaks (Supplementary figure 4A). In addition, an interaction between PGR and the histone acetyltransferase CBP/p300 was illustrated in hormone-stimulated granulosa cells (Supplementary figure 4B). A functional relationship between PGR binding and chromatin accessibility was further investigated through ATAC-seq in granulosa cells isolated from PGRWT and PGRKO mice. Of a total of 73,764 identified ATAC peaks, 1,499 sites were found to have significantly altered peak intensity in the absence of PGR (Figure 5A); interestingly, only 6 of the 1,499 sites had increased ATAC peak intensity in PGRKO compared to PGRWT. Whereas PGR-chromatin binding has been shown to favour proximal promoter regions in granulosa cells, the PGR-dependent ATAC-seq peaks were more evenly distributed throughout all genomic features (Figure 5B). Through RNA-seq in granulosa cells from PGRWT and PGRKO mice 8 h post-hCG treatment, 2 h after the peak of PGR abundance, 286 PGR-dependent DEGs were identified (Supplementary table 2). Comparison of these PGR-regulated genes and PGRKO ATAC signal showed a consistent reduction in ATAC-seq intensity at promoters of genes with PGR-dependent expression (Spearman correlation coefficient = 0.5116, p-value < 0.0001) (Figure 5C). 28 DEGs showed a significant difference in ATAC-seq peak intensity as determined through DiffBind (Figure 5C, light green), indicating that promoter accessibility for these genes is significantly influenced by PGR. Examples of the difference in PGR influence on promoters are shown for *Gas7*, whose promoter was differentially accessible in PGRKO, and *Abhd2*, the promoter accessibility of which was not significantly altered in the absence of PGR (Figure 5D).

**Figure 5:**
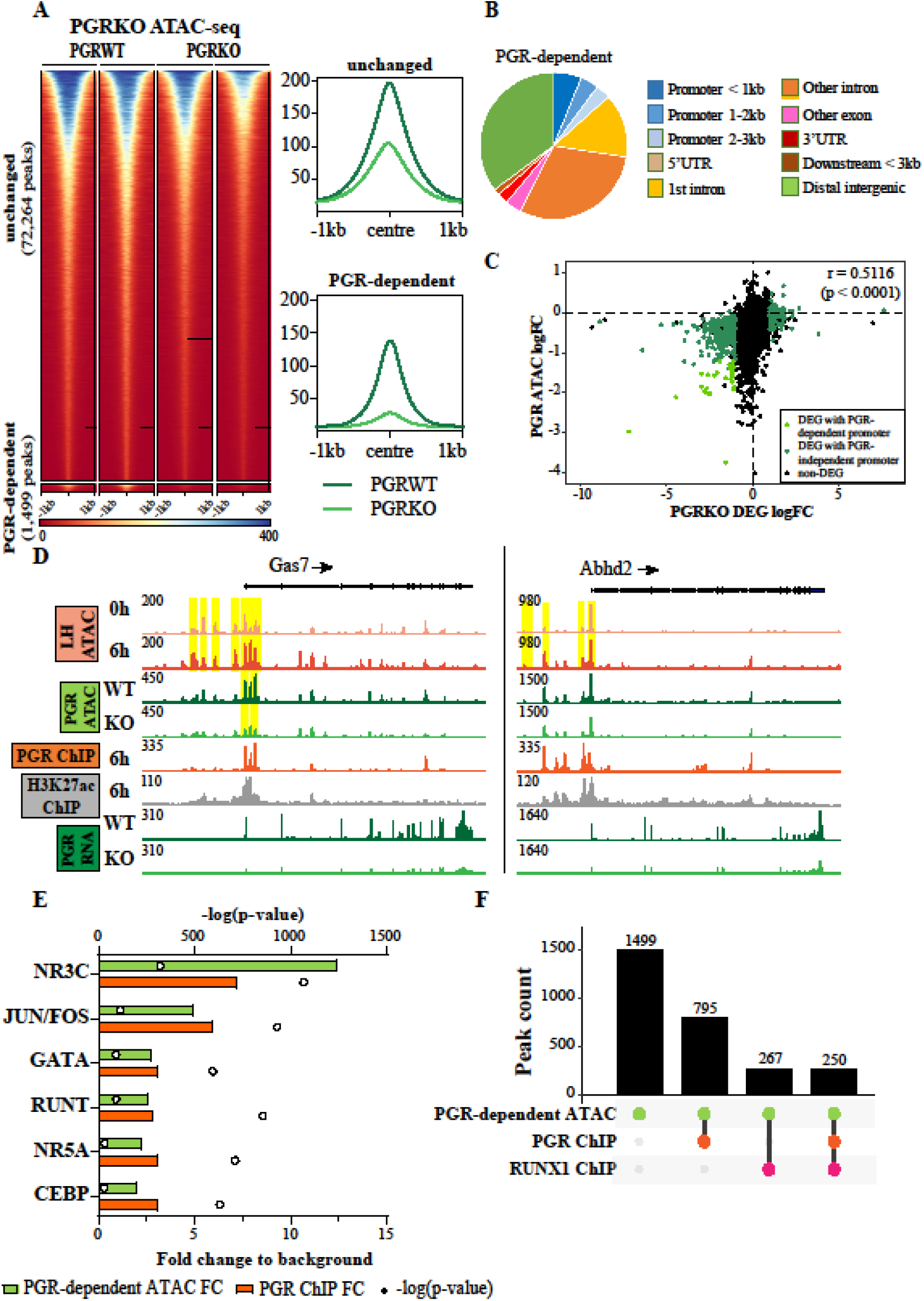
PGR role in mediating chromatin accessibility in granulosa cells. (A) Heatmap of PGR ATAC-peaks (left) that were unchanged (top) or differentially accessible between PGRKO WT and KO (bottom). Representative profile plots of each group are on the right. (B) Genomic distribution of peaks that were differentially enriched in PGRKO. (C) Correlation between expression level and promoter accessibility at PGR DEGs, shown as PGR DEG fold change vs ATAC fold change at their respective promoter. DEGs with PGR-dependent promoter accessibility are labelled light green, DEGs with promoter accessibility unchanged by PGR are labelled dark green. Correlation between RNA-seq and ATAC-seq intensity is determined through Spearman’s correlation coefficient (r = 0.5116, p < 0.0001) (D) Examples of LH ATAC-seq (track 1, 2), PGR ATAC-seq (track 3, 4), PGR and H3K27ac ChIP-seq (track 5, 6) and PGR RNA-seq (track 7, 8) signals at *Gas7* and *Abhd2*. ATAC peaks at promoters are highlighted in yellow. (E) Top most common known sequence motifs found to be enriched at PGR-dependent ATAC peaks (green), with the level of enrichment for the same motif at PGR binding sites as reference (orange). Bars indicate fold enrichment of motif to background (bottom x axis). Circles indicate −log(p-value) (top x axis). (F) Upset plot showing the number of overlapped peaks between PGR-dependent ATAC sites, PGR and RUNX1 binding sites.

Transcription factor motif enrichment analysis was performed through HOMER on PGR-driven ATAC peaks and compared with motifs enriched in PGR ChIP-seq. As with ChIP-seq, the most highly enriched and most significant motif identified in PGR-driven ATAC peak sequences was the canonical NR3C motif (12.4-fold over background). Again, several non-canonical motifs were also significantly enriched in ATAC-seq as with PGR ChIP-seq (Figure 5E). To examine the possibility that differential chromatin remodelling in PGRKO was mainly at sites with direct PGR-NR3C motif binding, we compared PGR and RUNX1 ChIP-seq cistromes with PGR-driven ATAC sites. Interestingly, 53% (795 of 1,499) of PGR-dependent ATAC sites were bound by PGR but only 17.8% (267 sites) were bound by RUNX1, almost all of which (250/267 sites) were PGR/RUNX1 co-bound (Figure 5F). This indeed suggests that PGR-dependent accessible chromatin sites were mainly bound by only PGR and not RUNX1.

### PGR-A is the key mediator of RUNX1 interaction and ovulatory gene regulation

We confirmed that PGR-A was the most important isoform in ovulation by verifying the disrupted ovulatory phenotype in both total PGRKO and AKO mice but not BKO mice, as previously documented ^23,24^ (Supplementary figure 5A). RNA-seq in granulosa cells obtained from PGRKO, AKO or BKO and WT littermates from each strain was analysed and hierarchical clustering analysis showed that the global gene expression pattern in AKO granulosa cells resembled that of total PGRKO, while BKO more closely resembled the WTs of each strain (Figure 6A). The loss of PGR-A resulted in the differential expression of 310 genes, 153 (or 49.4%) of which were also identified in the absence of both PGR isoforms (Figure 6B, Supplementary table 2). Among these, 48.6% and 58.5% of AKO and PGRKO DEG respectively were also differentially expressed in response to the LH surge. The loss of PGR-B, on the other hand, resulted in no significant transcriptional changes (Figure 6B), confirming that PGR-A plays a predominant role in the overall effect of PGR during ovulation. Examples of RNA-seq signals can be observed in PGR target genes such as *Pparg* and *Stard5* (Supplementary figure 5B, C), which exhibits dependence on PGR-A but not PGR-B.

**Figure 6:**
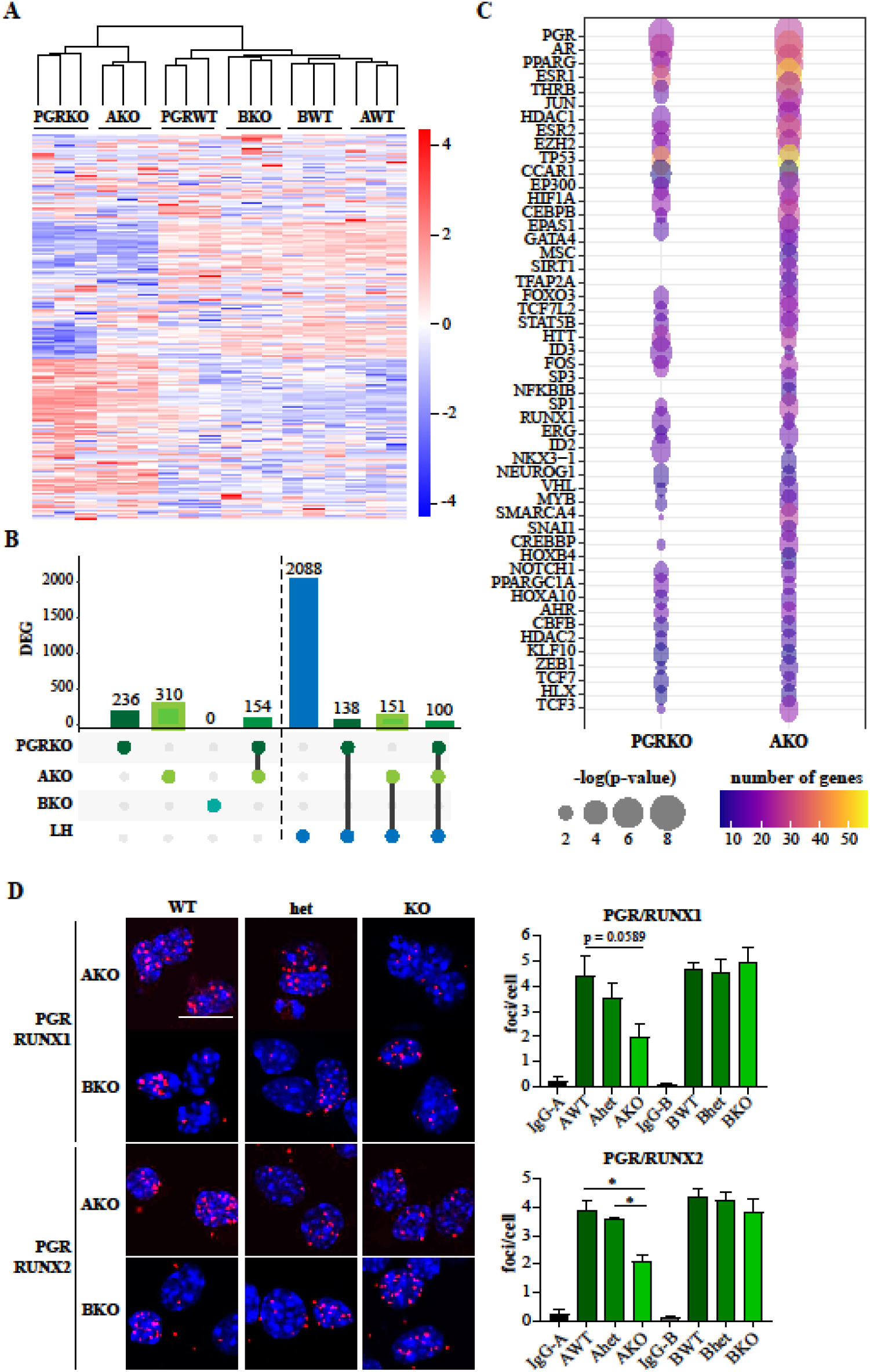
PGR isoform-driven transcriptome in peri-ovulatory granulosa cells through interaction with specific transcription factor partners. (A) Heatmap showing changes in gene expression in granulosa cells that are WT or KO for total PGR, PGR-A or PGR-B, each shown with the collective PGRKO, AKO and BKO DEGs. (B) Upset plot showing the number of overlapped genes between PGRKO/AKO/BKO DEG and LH DEG. (C) Top 50 upstream regulators of PGRKO and AKO DEGs as identified through IPA. Circle colour indicates the number of downstream DEG regulated by the upstream regulator, circle size indicates −log(p-value) of regulator enrichment. (D) PLA of PGR/RUNX1 (top) and PGR/RUNX2 (bottom) in *in vitro* WT/het/KO granulosa cells from AKO or BKO mice, treated for 6 h with hCG and R5020. Red indicates protein-protein interaction. Blue is DAPI nuclear stain. Scale bar = 10 μm. Quantification of PLA signal is displayed as the number of nuclear foci per cell, displayed as mean ± SEM, * = p-value < 0.05, N = 3 biological replicates (2 mice per replicate per genotype).

Upstream analysis of the DEG datasets using IPA analysis predicted a number of transcription factors as potential upstream regulators of the PGRKO and AKO responses (Figure 6C). As expected, PGR was among the most significant predicted upstream regulators, as well as other transcription factors we and others have shown to be targets of PGR action, such as PPARG ^33^ and HIF1A ^34^. Importantly, this independent approach also implicates RUNX1, CBFβ, JUN, FOS and CBP/p300 as mediators of the PGR-dependent transcriptional response. The lack of PGR-B effect on transcription regulation is suggestive of PGR isoform-specific interactomes in granulosa cells. To confirm whether the direct PGR interaction with RUNX was specific for PGR-A, we performed PLA for PGR/RUNX1 and PGR/RUNX2 in hCG/progestin-stimulated granulosa cells obtained from WT, het and KO animals from the AKO and BKO strains. A decrease in protein-protein interaction with both RUNX1 and RUNX2 was observed exclusively in AKO but not BKO granulosa cells (Figure 6D).

## 3. DISCUSSION

Safer, more accessible contraception as well as new treatments for anovulatory infertility remain key unmet needs in fertility management. In the era of high specificity, low side-effect drug development, this can be delivered once the molecular mechanisms driving the biological process are fully elaborated and validated. In this study we applied unbiased genome-wide approaches to characterise the global chromatin remodelling in granulosa cells responding to the *in vivo* ovulatory stimulus and identified the cell-specific gene regulatory network controlled by the action of PGR, the critical transcription factor required for ovulation (Figure 7). Our data confirm a specific role for PGR-A in granulosa cells through a unique interaction with RUNX transcription factors at non-canonical DNA motifs. We verified the cooperative binding of PGR with RUNX1/2 by demonstrating substantial overlap between PGR and RUNX1 cistromes as well as showing LH-dependent formation of nuclear complexes containing PGR and RUNX1/2 that depend on the presence of PGR-A but not PGR-B. Together, these findings support our hypothesis that the unique ovarian action of PGR involves the formation of PGR-A/RUNX transcriptional complexes with specific DNA sequence binding activity.

**Figure 7:**
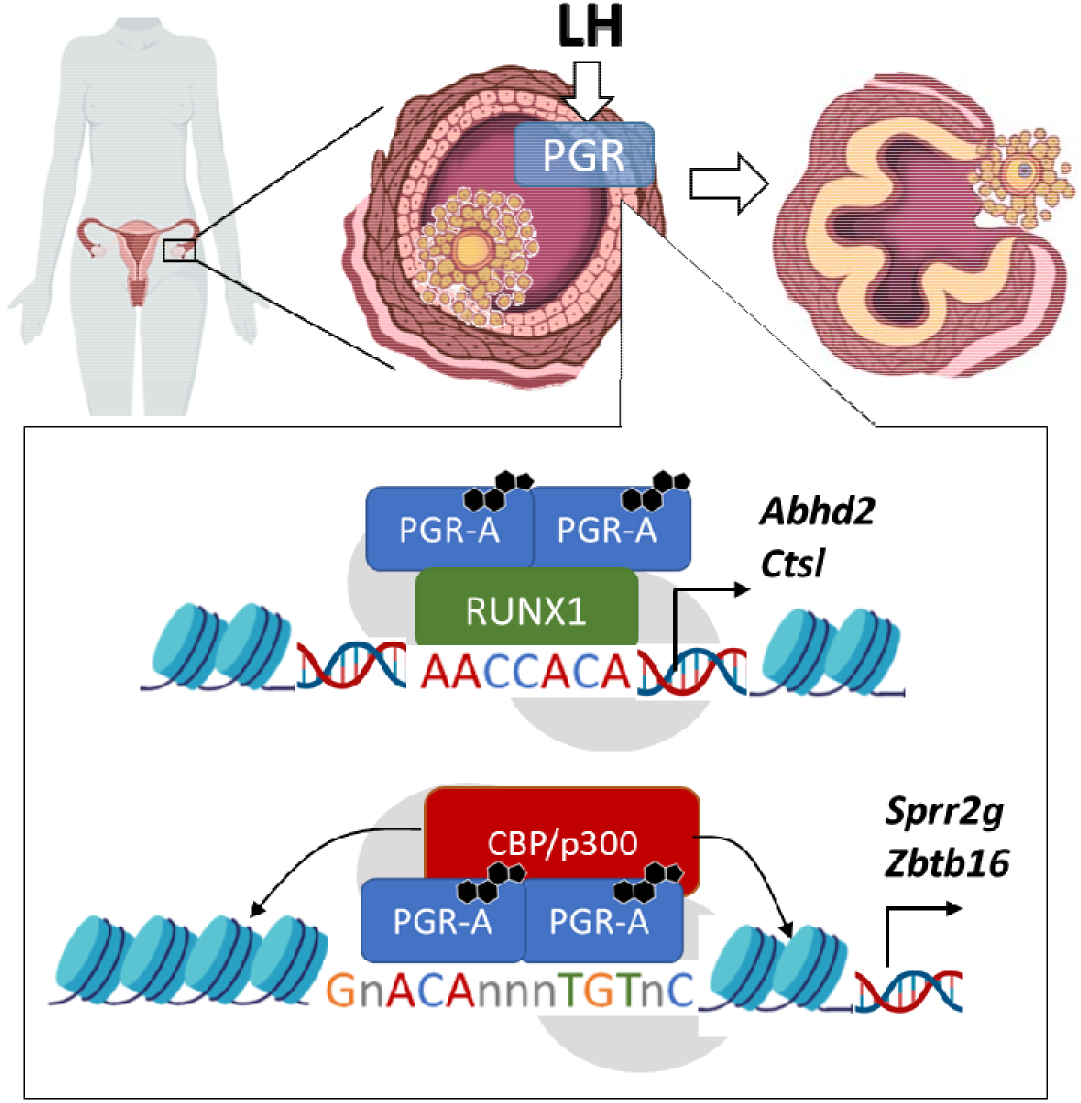
Model of PGR action in peri-ovulatory granulosa cells.

The LH surge leads to critical shifts in the chromatin landscape and the ovarian gene expression profile, eventually resulting in the activation of biological pathways that are important for follicle rupture. Markedly increased chromatin accessibility around promoters of ovulatory genes and also at many intergenic regions indicates global epigenetic changes in histone modifications throughout the genome within 6 h of the ovulatory stimulus, leading to the rapid and pervasive reprogramming of gene expression. Our approach also allowed comprehensive identification of the transcription factors involved. *De novo* motif identification and footprinting analysis confirm RUNX and JUN/FOS as well as NR5A and CEBP transcription factor binding at open chromatin sites. This is consistent with the reported induced expression of JUN and FOS ^35^, RUNX1/2 ^32,36^ and CEBPβ ^37^ by ovulatory stimulus, as well as an activation of the PKA-CREB1 signalling pathway ^38,39^.

Surprisingly, the canonical sequence motif used by NR3C steroid receptors, including PGR, shows the largest repression in chromatin accessibility in response to the ovulatory stimulus. Considering that PGR expression is highly induced in the peri-ovulatory period ^22^ and that it is a key determinant of ovulation ^6,40^, the loss of NR3C response element representation in open chromatin regions suggests that PGR-chromatin interaction is less dependent on this consensus sequence and supports our previous report that PGR binding sites in granulosa cells are exceptionally enriched in RUNX and JUN/FOS sequence motifs ^22^. Tissue-specific PGR interactomes have been shown to be important for the diversity of PGR transactivation functions in various contexts ^41^. Here, an interaction between PGR and RUNX1 is described for the first time, in which PGR and RUNX1/2 form a very close physical association specifically within the nucleus of granulosa cells in response to ovulatory cues, strongly suggesting that they reside in a cooperative transcriptional complex. Further evidence that PGR/RUNX1 interact in a transcriptional complex comes from the remarkable overlap in PGR/RUNX1 co-occupancy on granulosa cell chromatin (70% of PGR binding sites overlapping with RUNX1), especially at ovulatory genes. The very high prevalence of promoter-centric RUNX1 binding and RUNX1/PGR co-binding mimics the promoter-centric enrichment of the PGR cistrome we previously reported to be highly specific in granulosa cells and not other PGR target tissues ^12,22^. We now show that this selective PGR-promoter interaction is dependent on RUNX1 and is also not reciprocal, since RUNX1 preferably targets proximal promoters both in the presence or absence of PGR. The predominance of PGR and RUNX1 binding identified at LH-regulated genes is equally striking. Together the results imply that RUNX1 directs PGR binding to promoter regions and they cooperatively regulate a significant proportion of the ovulatory gene expression profile. Consistent with this conclusion, RUNX1 has been previously reported to behave as a pioneer factor for the SWI/SNF chromatin remodelling complex ^42^ and has a role in programming chromatin conformation ^43^. Reports of the interaction between members of the RUNX family and NR3C members, AR and GR, leading either to the repression ^44^ or transactivation of these transcription factors ^44–47^, as well as RUNX1 tethering effect on ERα^48^, further substantiate the conclusion that RUNX transcription factors tether PGR to granulosa-specific chromatin targets. Utilising isoform-specific knockout mouse models, we also show that PGR-A is the predominant isoform obligatory for ovulation. The unique granulosa cell function of PGR-A is likely due to isoform-specific interaction with other transcription mediators, exemplified by the specific interaction between PGR-A and RUNX1. PGR is also known to have an active role in mediating chromatin accessibility through the recruitment of chromatin remodellers ^14,15^. The interaction between PGR and the histone acetyltransferase CBP/p300, identified through PLA, and altered accessibility at 1,499 chromatin sites in PGRKO granulosa cells shows that PGR is involved in chromatin reprogramming in ovulation. Together this indicates that tissue-specific PGR response is driven by proximal promoter-centric PGR/RUNX interaction as well as RUNX1-independent PGR binding outside of proximal regulatory regions that coordinately facilitates chromatin remodelling.

The physical interaction of PGR with other transcription factors, such as NR5A2 (LRH1) and JUND, is also identified. Although a low level of PLA signal suggests that PGR and LRH1 are closely co-localised, a functional interaction is less likely as they possess different windows of expression in granulosa cells (PGR is completely absent in granulosa cells prior to the LH surge whereas LRH1 is expressed from primordial stage up to corpus luteum ^49^. In support of this, less than 20% of PGR-regulated genes were also found to be LRH1 targets in peri-ovulatory granulosa cells ^2^. On the other hand, JUN/FOS transcription factors are highly induced by the LH surge in granulosa cells, similar to PGR ^35,50^. PGR/JUND PLA signal was abundant and substantial direct occupancy of the JUN/FOS motif was exclusive to LH-induced open chromatin sites. We suggest that JUN/FOS activation and chromatin binding is a parallel pathway to PGR/RUNX1/2 complexes and these transcription factors cooperatively mediate transcriptional activation of ovulatory genes. Further studies of JUN/FOS cistromes and transcriptomes in peri-ovulatory granulosa cells are necessary to define their contribution to PGR action and ovulatory function.

In summary, our results have clearly defined a unique mechanistic model of PGR action in granulosa cells which involves PGR-A interaction with RUNX1 to drive the specialised ovulatory PGR-mediated gene regulatory network. This ovarian-specific molecular mechanism of PGR action is distinct from the mechanisms in other PGR-responsive organs. These new insights are critical for understanding and addressing causes of anovulatory infertility and for the development of ovulation-specific contraceptives with minimal side-effects that often arise from progestin therapies.

## 4. METHODS

### Reagents and antibodies

Unless otherwise stated, reagents were purchased from Sigma-Aldrich (St. Louis, MO, USA).

### Animals

21-day old CBA x C57BL/6 F1 (CBAF1) mice were obtained from the Laboratory Animal Services (University of Adelaide). Null mutant mouse strains with ablation of either all PGR isoforms, or specific ablation of PGR-A or PGR-B ^7,23,24^ were obtained from the Jackson Laboratory (Bar Harbor, USA). PGRKO mice are a targeted mutation strain with the Jackson Laboratory designation Pgr^tm1Bwo^. PGR-A KO (AKO) mice are a targeted mutation strain with the Jackson Laboratory designation Pgr^tm1Omc^. PGR-B KO (BKO) mice are a targeted mutation strain with the Jackson Laboratory designation Pgr^tm2Omc^. Mice from KO strains are routinely genotyped from ear or tail biopsies before allocation to experiments and confirmed from replicate biopsies collected at the time of experiments. All mice were maintained in 12 h light / 12 h dark conditions and given water and rodent chow *ad libitum*. Littermate females of each genotype were allocated for experiments at 21 days of age. All experiments were approved by The University of Adelaide Animal Ethics Committee and were conducted in accordance with the Australian Code of Practice for the Care and Use of Animals for Scientific Purposes (ethics number m/2015/075, m/2018/100, m/2018/122, m/2018/117).

### Granulosa cell culture and hormone treatment

CBAF1 female mice were stimulated with 5 IU eCG (Lee BioSolutions, Maryland Heights, USA) and culled at 44 hours post-eCG, upon which ovaries were dissected and granulosa cells isolated. Cumulus-oocyte complexes were removed and granulosa cells were counted before being seeded into an 8-well fibronectin-coated chamber slide (minimum 100,000 cells/well). Cells were cultured in DMEM:F12 media (Thermo Fisher) at 37°C, 5% CO_2_ for 90 minutes then washed with PBS to remove debris. Cells were incubated overnight and were treated with 2 IU/ml hCG (Merck Sharp & Dohme B.V., Haarlem, Netherlands) and 100 nM R5020 (PerkinElmer, Waltham, USA) for indicated durations at 37°C, 5% CO_2_.

### ATAC-seq

ATAC-seq was performed as previously described ^51^. Briefly, granulosa cells were collected from superovulated female CBAF1 or PGRKO mice at 0 h or 6 h post-hCG as above. Cells from 2-3 animals were pooled together for one biological replicate, two replicates were obtained per condition. Granulosa cells were lysed and tagmentation was performed on the nuclei using the Illumina Tagment DNA TDE1 Enzyme and Buffer Kits (Illumina, San Diego, USA). DNA purification was performed using the Monarch PCR & DNA Cleanup Kit (NEB, Ipswich, USA). Libraries were then amplified and purified before being size selected using magnetic AMPure XP Beads (Beckman Coulter, Brea, USA). Library fragmentation was confirmed using the LabChIP GX Bioanalyser (PerkinElmer) and sequencing was performed on the HiSeq2500 Sequencing System (Illumina). For all datasets, adapters were trimmed from 150-base sequences using Cutadapt ^52^ and aligned to the mm10 mouse genome using Bowtie2 algorithm ^53^. Alignment was filtered using samtools for non-mitochondrial reads with MAPQ ≥ 30 cut-off and was de-duplicated using Picard tools. Peak calling from read count followed the algorithm for Model-based Analysis for ChIP-Seq (MACS2) with a q-value cut-off = 10^−10^ and a mouse genome size of 1.87×10^9 54^. Differential binding analysis for LH ATAC-seq and PGR ATAC-seq was performed using DiffBind ^55^ with DESeq2 linear model, with differentially enriched sites determined to have FDR ≤ 0.05. Genomic distribution of peak subsets were determined using the ChIPseeker package ^56^. Gene ontology enrichment analysis was performed using GREAT ^57^. Differential enrichment of transcription factor binding motif and DNA footprinting analysis was performed using TOBIAS ^58^. Motif analysis for *de novo* sequence motifs was performed using HOMER motif finding algorithm with random 200 bp-long sequences from the mouse genome used to estimate motif frequency in random sequence ^59^. Visual representation of ATAC signal was through the UCSC Genome Browser (Santa Cruz, USA).

### RNA-seq

Three-week old female mice, either CBAF1 (for LH RNA-seq) or WT and KO from PGRKO, AKO and BKO strains (for PGR RNA-seq) were hormonally stimulated as above, before being culled at 8 h post-hCG injection for granulosa cell collection. For LH RNA-seq, eCG-treated mice that did not receive hCG injection served as negative control. For each condition, four individual replicates were collected, each containing pooled cells from three animals. RNA was extracted from granulosa cells using RNeasy Mini kit (Qiagen, Chadstone, VIC, Australia). RNA quality was assessed using the RNA ScreenTape System (Agilent, Santa Clara, CA, USA).

For LH response RNA-seq, RNA was subjected to rRNA depletion using a modified Ribozero method and library preparation using Clontech SMARTer Stranded RNA-Seq Kits (Takara Bio, Shiga, Japan). Sequencing was performed on the Illumina HiSeq2500 100bp PE Rapid run (Ramaciotti Centre for Genomics, Kensington, Australia). For PGR RNA-seq, the library was prepared using the Universal RNA-Seq library kit with NuQuant Mouse AnyDeplete (Nugen, Redwood City, USA) and sequencing was performed on the NovaSeq 6000 S1 Sequencing System (South Australian Health & Medical Research Institute, Adelaide, Australia). Overrepresented adapters were checked using FastQC (https://www.bioinformatics.babraham.ac.uk/projects/fastqc/) and adapters were trimmed using AdapterRemoval ^60^ when required. Quasi-aligment of each dataset to the GENCODE mouse transcriptome (GRCm38.p6, M25 release) and transcript quantification were through Salmon ^61^. Expression profiles and differential gene expression were assessed with limma ^62^ and edgeR ^63^ using mean-variance estimates from log counts as described in the limma voom method. Differential gene expression for each experiment was defined as a fold-change ≥ 2 (|log FC| ≥ 1) and Benjamini-Hochberg adjusted p-value ≤ 0.01. Upstream regulator analysis of DEGs was through IPA software (QIAGEN). For visualisation of RNA-seq on the UCSC Genome Browser, read coverage was processed using deepTools ^64^.

### ChIP-seq

Granulosa cells from CBAF1 female mice were collected by puncturing ovarian follicles 44 h after eCG (0 h hCG) and 6 h after hCG stimulation. Two biological replicates were obtained for each sample from 10 mice, each with at least 1×10^7^ cells. ChIP-seq for RUNX1 was performed by Active Motif (Carlsbad, USA) as previously described using in-house RUNX1 antibody ^65^. For all datasets, sequences were aligned to the mm10 mouse genome using Bowtie2 algorithm. Peak calling from read count followed the algorithm for MACS2 with a p-value cut-off = 10^−10^. Overlapped peaks determined using ChIPseeker were used as the consensus data in all subsequent comparisons. Differential binding analysis for RUNX1 ChIP-seq was performed using DiffBind as described above. Peak characterisation was determined using the ChIPseeker package. Motif analysis for known and *de novo* sequence motifs was performed using HOMER as described above. Visualisation of ChIP-seq signal was through the UCSC Genome Browser.

### Proximity ligation assay (PLA)

PLA was performed on cultured granulosa cells using the Duolink PLA Probes and PLA Fluorescence in situ Detection Kit Red (Sigma) following the manufacturer’s protocol. Briefly, cultured granulosa cells were fixed in 4% paraformaldehyde and permeabilised with PBS + 0.01% Triton X-100 for 1 hour at room temperature. Cells were blocked with Blocking Buffer for 1 hour at 37°C and incubated with primary antibody couples (PGR – Cell Signalling Technology (CST); RUNX1 – Santa Cruz Biotechnology (SCB); RUNX2 – SCB; c-JUN – SCB; JUNB – SCB; JUND – SCB; LRH1 – Perseus Protemics, CBP/p300 – CST) diluted in Antibody Diluent for 2 hours at room temperature or overnight at 4°C. Then, cells were incubated with PLA probes of appropriate species for 1 hour at 37°C, oligo probes were ligated for 30 minutes at 37°C and the amplification reaction was at 37°C for a minimum of 100 minutes. Between steps, cells were washed using the provided wash buffers. Slides were mounted with Prolong Gold Mounting Media with DAPI (Thermo Fisher), cured for at least 1 hour in the dark and were stored at −20°C prior to imaging by Olympus confocal microscope. The nuclear boundary for each cell was determined through DAPI staining. PLA signals were identified as fluorescent puncta in the nucleus and quantified using the ‘Count Maxima’ function in ImageJ for each cellular compartment. Significant differences between time points were determined through one-way ANOVA with Tukey test for multiple comparison.

### Data availability

The datasets generated and/or analysed during the current study are available in the NCBI Gene Expression Omnibus repository: GSE166878 (LH and PGRKO ATAC-seq), GSE178314 (LH RNA-seq), GSE168213 (PGRKO RNA-seq), GSE152941 (RUNX1 ChIP-seq).

## Acknowledgement

The RUNX1 antibody for ChIP-seq was a kind gift from Dr Yoram Groner and Dr Ditsa Levanon, the Weizmann Institute of Science, Israel. This work was partially supported by the Intramural Research Program of National Institute of Environmental Health Sciences in the USA (Z01-ES102965 for HHCY).

## Author contribution

D.L.R conceived and designed the study. T.D.D, J.B, R.L.R and D.L.R conceived and designed the experiments. T.D.D, B.N, K.M.S, M.N, A.E, Y.Y.W performed experiments. T.D.D and J.B analysed data. T.D.D, S.C.B, H.H.C.Y, R.L.R and D.L.R interpreted results. T.D.D prepared figures. T.D.D and D.L.R drafted manuscript. All authors contributed to manuscript editing and revision.

## REFERENCES

1 Russell, D. L. & Robker, R. L. in The Ovary (Third Edition) (eds Peter C. K. Leung & Eli Y. Adashi) 217–234 (Academic Press, 2019).

2 Hughes, C. H. K. & Murphy, B. D. Nuclear receptors: Key regulators of somatic cell functions in the ovulatory process. Molecular Aspects of Medicine, 100937, doi:https://doi.org/10.1016/j.mam.2020.100937 (2020).

3 Wu, S.-P., Li, R. & DeMayo, F. J. Progesterone Receptor Regulation of Uterine Adaptation for Pregnancy. Trends in Endocrinology & Metabolism 29, 481–491, doi:https://doi.org/10.1016/j.tem.2018.04.001 (2018).

4 Conneely, O. M., Mulac-Jericevic, B. & Arnett-Mansfield, R. in Progestins and the Mammary Gland. (eds O. Conneely & C. Otto) 175–185 (Springer Berlin Heidelberg).

5 Robker, R. L., Russell, D. L., Espey, L. L., Lydon, J. P., O’Malley, B. W. & Richards, J. S. Progesterone-regulated genes in the ovulation process: ADAMTS-1 and cathepsin L proteases. Proc Natl Acad Sci U S A 97, 4689–4694 (2000).

6 Park, C. J., Lin, P.-C., Zhou, S., Barakat, R., Bashir, S. T., Choi, J. M., Cacioppo, J. A., Oakley, O. R., Duffy, D. M., Lydon, J. P. & Ko, C. J. Progesterone Receptor Serves the Ovary as a Trigger of Ovulation and a Terminator of Inflammation. Cell Reports 31, 107496, doi:https://doi.org/10.1016/j.celrep.2020.03.060 (2020).

7 Lydon, J. P., DeMayo, F. J., Funk, C. R., Mani, S. K., Hughes, A. R., Montgomery, C. A., Jr., Shyamala, G., Conneely, O. M. & O’Malley, B. W. Mice lacking progesterone receptor exhibit pleiotropic reproductive abnormalities. Genes Dev 9, 2266–2278 (1995).

8 Ledger, W. L., Sweeting, V. M., Hillier, H. & Baird, D. T. Inhibition of ovulation by low-dose mifepristone (RU 486). Human Reproduction 7, 945–950, doi:10.1093/oxfordjournals.humrep.a137776 (1992).

9 Brache, V., Cochon, L., Jesam, C., Maldonado, R., Salvatierra, A. M., Levy, D. P., Gainer, E. & Croxatto, H. B. Immediate pre-ovulatory administration of 30 mg ulipristal acetate significantly delays follicular rupture. Human Reproduction 25, 2256–2263, doi:10.1093/humrep/deq157 (2010).

10 Clemens, J. W., Robker, R. L., Kraus, W. L., Katzenellenbogen, B. S. & Richards, J. S. Hormone induction of progesterone receptor (PR) messenger ribonucleic acid and activation of PR promoter regions in ovarian granulosa cells: evidence for a role of cyclic adenosine 3’,5’-monophosphate but not estradiol. Mol Endocrinol 12, 1201–1214, doi:10.1210/mend.12.8.0157 (1998).

11 Ismail, P. M., Li, J., DeMayo, F. J., O’Malley, B. W. & Lydon, J. P. A Novel LacZ Reporter Mouse Reveals Complex Regulation of the Progesterone Receptor Promoter During Mammary Gland Development. Molecular Endocrinology 16, 2475–2489, doi:10.1210/me.2002-0169 (2002).

12 Yin, P., Roqueiro, D., Huang, L., Owen, J. K., Xie, A., Navarro, A., Monsivais, D., Coon V, J. S., Kim, J. J., Dai, Y. & Bulun, S. E. Genome-Wide Progesterone Receptor Binding: Cell Type-Specific and Shared Mechanisms in T47D Breast Cancer Cells and Primary Leiomyoma Cells. PLOS ONE 7, e29021, doi:10.1371/journal.pone.0029021 (2012).

13 Li, X. & O’Malley, B. W. Unfolding the Action of Progesterone Receptors. Journal of Biological Chemistry 278, 39261–39264, doi:10.1074/jbc.R300024200 (2003).

14 Han, S. J., DeMayo, F. J., Xu, J., Tsai, S. Y., Tsai, M.-J. & O’Malley, B. W. Steroid Receptor Coactivator (SRC)-1 and SRC-3 Differentially Modulate Tissue-Specific Activation Functions of the Progesterone Receptor. Molecular Endocrinology 20, 45–55, doi:10.1210/me.2005-0310 (2006).

15 Béguelin, W., Díaz Flaqué, M. C., Proietti, C. J., Cayrol, F., Rivas, M. A., Tkach, M., Rosemblit, C., Tocci, J. M., Charreau, E. H., Schillaci, R. & Elizalde, P. V. Progesterone receptor induces ErbB-2 nuclear translocation to promote breast cancer growth via a novel transcriptional effect: ErbB-2 function as a coactivator of Stat3. Molecular and cellular biology 30, 5456–5472, doi:10.1128/MCB.00012-10 (2010).

16 Malbeteau, L., Poulard, C., Languilaire, C., Mikaelian, I., Flamant, F., Le Romancer, M. & Corbo, L. PRMT1 Is Critical for the Transcriptional Activity and the Stability of the Progesterone Receptor. iScience 23, 101236, doi:https://doi.org/10.1016/j.isci.2020.101236 (2020).

17 Cho, H., Orphanides, G., Sun, X., Yang, X. J., Ogryzko, V., Lees, E., Nakatani, Y. & Reinberg, D. A human RNA polymerase II complex containing factors that modify chromatin structure. Molecular and cellular biology 18, 5355–5363, doi:10.1128/mcb.18.9.5355 (1998).

18 Dong, X., Challis, J. R. & Lye, S. J. Intramolecular interactions between the AF3 domain and the C-terminus of the human progesterone receptor are mediated through two LXXLL motifs. J Mol Endocrinol 32, 843–857, doi:10.1677/jme.0.0320843 (2004).

19 Merlino, A. A., Welsh, T. N., Tan, H., Yi, L. J., Cannon, V., Mercer, B. M. & Mesiano, S. Nuclear Progesterone Receptors in the Human Pregnancy Myometrium: Evidence that Parturition Involves Functional Progesterone Withdrawal Mediated by Increased Expression of Progesterone Receptor-A. The Journal of Clinical Endocrinology & Metabolism 92, 1927–1933, doi:10.1210/jc.2007-0077 (2007).

20 Menendez, J. A., Peirce, S. K., Papadimitropoulou, A., Cuyàs, E., Steen, T. V., Verdura, S., Vellon, L., Chen, W. Y. & Lupu, R. Progesterone receptor isoform-dependent cross-talk between prolactin and fatty acid synthase in breast cancer. Aging (Albany NY) 12, 24671–24692, doi:10.18632/aging.202289 (2020).

21 Teilmann, S. C., Clement, C. A., Thorup, J., Byskov, A. G. & Christensen, S. T. Expression and localization of the progesterone receptor in mouse and human reproductive organs. J Endocrinol 191, 525–535, doi:10.1677/joe.1.06565 (2006).

22 Dinh, D. T., Breen, J., Akison, L. K., DeMayo, F. J., Brown, H. M., Robker, R. L. & Russell, D. L. Tissue-specific progesterone receptor-chromatin binding and the regulation of progesterone-dependent gene expression. Scientific reports 9, 11966–11966, doi:10.1038/s41598-019-48333-8 (2019).

23 Mulac-Jericevic, B., Mullinax, R. A., DeMayo, F. J., Lydon, J. P. & Conneely, O. M. Subgroup of Reproductive Functions of Progesterone Mediated by Progesterone Receptor-B Isoform. Science 289, 1751–1754, doi:10.1126/science.289.5485.1751 (2000).

24 Mulac-Jericevic, B., Lydon, J. P., DeMayo, F. J. & Conneely, O. M. Defective mammary gland morphogenesis in mice lacking the progesterone receptor B isoform. Proceedings of the National Academy of Sciences 100, 9744–9749, doi:10.1073/pnas.1732707100 (2003).

25 Hu, M., Li, J., Zhang, Y., Li, X., Brännström, M., Shao, L. R. & Billig, H. Endometrial progesterone receptor isoforms in women with polycystic ovary syndrome. Am J Transl Res 10, 2696–2705 (2018).

26 Wetendorf, M., Li, R., Wu, S.-P., Liu, J., Creighton, C. J., Wang, T., Janardhan, K. S., Willson, C. J., Lanz, R. B., Murphy, B. D., Lydon, J. P. & DeMayo, F. J. Constitutive expression of progesterone receptor isoforms promotes the development of hormone-dependent ovarian neoplasms. Science Signaling 13, eaaz9646, doi:10.1126/scisignal.aaz9646 (2020).

27 Liu, J., Park, E.-S. & Jo, M. Runt-related transcription factor 1 regulates luteinized hormone-induced prostaglandin-endoperoxide synthase 2 expression in rat periovulatory granulosa cells. Endocrinology 150, 3291–3300 (2009).

28 Liu, J., Park, E.-S., Curry, T. E., Jr. & Jo, M. Periovulatory expression of hyaluronan and proteoglycan link protein 1 (Hapln1) in the rat ovary: hormonal regulation and potential function. Mol Endocrinol 24, 1203–1217, doi:10.1210/me.2009-0325 (2010).

29 Lee-Thacker, S., Choi, Y., Taniuchi, I., Takarada, T., Yoneda, Y., Ko, C. & Jo, M. Core binding factor β expression in ovarian granulosa cells is essential for female fertility. Endocrinology 159, 2094–2109 (2018).

30 Lee-Thacker, S., Jeon, H., Choi, Y., Taniuchi, I., Takarada, T., Yoneda, Y., Ko, C. & Jo, M. Core Binding Factors are essential for ovulation, luteinization, and female fertility in mice. Scientific Reports 10, 9921, doi:10.1038/s41598-020-64257-0 (2020).

31 Nicol, B., Grimm, S. A., Chalmel, F., Lecluze, E., Pannetier, M., Pailhoux, E., Dupin-De-Beyssat, E., Guiguen, Y., Capel, B. & Yao, H. H. C. RUNX1 maintains the identity of the fetal ovary through an interplay with FOXL2. Nature communications 10, 5116 (2019). <https://doi.org/10.1038/s41467-019-13060-1>.

32 Park, E.-S., Park, J., Franceschi, R. T. & Jo, M. The role for runt related transcription factor 2 (RUNX2) as a transcriptional repressor in luteinizing granulosa cells. Molecular and Cellular Endocrinology 362, 165–175, doi:https://doi.org/10.1016/j.mce.2012.06.005 (2012).

33 Kim, J., Sato, M., Li, Q., Lydon, J. P., DeMayo, F. J., Bagchi, I. C. & Bagchi, M. K. Peroxisome Proliferator-Activated Receptor γ Is a Target of Progesterone Regulation in the Preovulatory Follicles and Controls Ovulation in Mice. Molecular and cellular biology 28, 1770–1782, doi:10.1128/mcb.01556-07 (2008).

34 Kim, J., Bagchi, I. C. & Bagchi, M. K. Signaling by hypoxia-inducible factors is critical for ovulation in mice. Endocrinology 150, 3392–3400, doi:10.1210/en.2008-0948 (2009).

35 Sharma, S. C. & Richards, J. S. Regulation of AP1 (Jun/Fos) factor expression and activation in ovarian granulosa cells. Relation of JunD and Fra2 to terminal differentiation. J Biol Chem 275, 33718–33728, doi:10.1074/jbc.M003555200 (2000).

36 Jo, M. & Curry, T. E., Jr. Luteinizing Hormone-Induced RUNX1 Regulates the Expression of Genes in Granulosa Cells of Rat Periovulatory Follicles. Molecular Endocrinology 20, 2156–2172, doi:10.1210/me.2005-0512 (2006).

37 Fan, H.-Y., Liu, Z., Johnson, P. F. & Richards, J. S. CCAAT/Enhancer-Binding Proteins (C/EBP)-α and -β Are Essential for Ovulation, Luteinization, and the Expression of Key Target Genes. Molecular Endocrinology 25, 253–268, doi:10.1210/me.2010-0318 (2011).

38 Russell, D. L., Doyle, K. M. H., Gonzales-Robayna, I., Pipaon, C. & Richards, J. S. Egr-1 Induction in Rat Granulosa Cells by Follicle-Stimulating Hormone and Luteinizing Hormone: Combinatorial Regulation By Transcription Factors Cyclic Adenosine 3’,5’-Monophosphate Regulatory Element Binding Protein, Serum Response Factor, Sp1, and Early Growth Response Factor-1. Molecular Endocrinology 17, 520–533, doi:10.1210/me.2002-0066 (2003).

39 Puri, P., Little-Ihrig, L., Chandran, U., Law, N. C., Hunzicker-Dunn, M. & Zeleznik, A. J. Protein Kinase A: A Master Kinase of Granulosa Cell Differentiation. Scientific Reports 6, 28132, doi:10.1038/srep28132 (2016).

40 Robker, R. L., Russell, D. L., Espey, L. L., Lydon, J. P., O’Malley, B. W. & Richards, J. S. Progesterone-regulated genes in the ovulation process: ADAMTS-1 and cathepsin L proteases. Proceedings of the National Academy of Sciences 97, 4689–4694, doi:10.1073/pnas.080073497 (2000).

41 Grimm, S. L., Hartig, S. M. & Edwards, D. P. Progesterone Receptor Signaling Mechanisms. Journal of Molecular Biology 428, 3831–3849, doi:https://doi.org/10.1016/j.jmb.2016.06.020 (2016).

42 Chuang, L. S. H., Ito, K. & Ito, Y. RUNX family: Regulation and diversification of roles through interacting proteins. International Journal of Cancer 132, 1260–1271, doi:10.1002/ijc.27964 (2013).

43 Barutcu, A. R., Hong, D., Lajoie, B. R., McCord, R. P., van Wijnen, A. J., Lian, J. B., Stein, J. L., Dekker, J., Imbalzano, A. N. & Stein, G. S. RUNX1 contributes to higher-order chromatin organization and gene regulation in breast cancer cells. Biochimica et Biophysica Acta (BBA) - Lipids and Lipid Metabolism 1859, 1389–1397, doi:10.1016/j.bbagrm.2016.08.003 (2016).

44 Kawate, H., Wu, Y., Ohnaka, K. & Takayanagi, R. Mutual transactivational repression of Runx2 and the androgen receptor by an impairment of their normal compartmentalization. The Journal of Steroid Biochemistry and Molecular Biology 105, 46–56, doi:https://doi.org/10.1016/j.jsbmb.2006.11.020 (2007).

45 Yang, R., Browne, J. A., Eggener, S. E., Leir, S.-H. & Harris, A. A novel transcriptional network for the androgen receptor in human epididymis epithelial cells. Molecular Human Reproduction 24, 433–443, doi:10.1093/molehr/gay029 (2018).

46 Ning, Y.-M. & Robins, D. M. AML3/CBFα1 Is Required for Androgen-specific Activation of the Enhancer of the Mouse Sex-limited Protein (Slp) Gene. Journal of Biological Chemistry 274, 30624–30630, doi:10.1074/jbc.274.43.30624 (1999).

47 Takayama, K.-i., Suzuki, T., Tsutsumi, S., Fujimura, T., Urano, T., Takahashi, S., Homma, Y., Aburatani, H. & Inoue, S. RUNX1, an androgen- and EZH2-regulated gene, has differential roles in AR-dependent and -independent prostate cancer. Oncotarget 6 (2014).

48 Stender, J. D., Kim, K., Charn, T. H., Komm, B., Chang, K. C. N., Kraus, W. L., Benner, C., Glass, C. K. & Katzenellenbogen, B. S. Genome-Wide Analysis of Estrogen Receptor α DNA Binding and Tethering Mechanisms Identifies Runx1 as a Novel Tethering Factor in Receptor-Mediated Transcriptional Activation. Molecular and cellular biology 30, 3943–3955, doi:10.1128/mcb.00118-10 (2010).

49 Falender, A. E., Lanz, R., Malenfant, D., Belanger, L. & Richards, J. S. Differential Expression of Steroidogenic Factor-1 and FTF/LRH-1 in the Rodent Ovary. Endocrinology 144, 3598–3610, doi:10.1210/en.2002-0137 (2003).

50 Choi, Y., Rosewell, K. L., Brännström, M., Akin, J. W., Curry Jr, T. E. & Jo, M. FOS, a Critical Downstream Mediator of PGR and EGF Signaling Necessary for Ovulatory Prostaglandins in the Human Ovary. The Journal of Clinical Endocrinology & Metabolism 103, 4241–4252 (2018).

51 Corces, M. R., Trevino, A. E., Hamilton, E. G., Greenside, P. G., Sinnott-Armstrong, N. A., Vesuna, S., Satpathy, A. T., Rubin, A. J., Montine, K. S., Wu, B., Kathiria, A., Cho, S. W., Mumbach, M. R., Carter, A. C., Kasowski, M., Orloff, L. A., Risca, V. I., Kundaje, A., Khavari, P. A., Montine, T. J., Greenleaf, W. J. & Chang, H. Y. An improved ATAC-seq protocol reduces background and enables interrogation of frozen tissues. Nature Methods 14, 959–962, doi:10.1038/nmeth.4396 (2017).

52 Martin, M. Cutadapt removes adapter sequences from high-throughput sequencing reads. EMBnet. journal 17, 10–12 (2011).

53 Langmead, B. & Salzberg, S. L. Fast gapped-read alignment with Bowtie 2. Nature Methods 9, 357–359, doi:10.1038/nmeth.1923 (2012).

54 Zhang, Y., Liu, T., Meyer, C. A., Eeckhoute, J., Johnson, D. S., Bernstein, B. E., Nusbaum, C., Myers, R. M., Brown, M., Li, W. & Liu, X. S. Model-based analysis of ChIP-Seq (MACS). Genome Biol 9, R137, doi:10.1186/gb-2008-9-9-r137 (2008).

55 Stark R, B. G. DiffBind: differential binding analysis of ChIP-Seq peak data. (2011).

56 Yu, G., Wang, L. G. & He, Q. Y. ChIPseeker: an R/Bioconductor package for ChIP peak annotation, comparison and visualization. Bioinformatics 31, 2382–2383, doi:10.1093/bioinformatics/btv145 (2015).

57 McLean, C. Y., Bristor, D., Hiller, M., Clarke, S. L., Schaar, B. T., Lowe, C. B., Wenger, A. M. & Bejerano, G. GREAT improves functional interpretation of cis-regulatory regions. Nature biotechnology 28, 495 (2010).

58 Bentsen, M., Goymann, P., Schultheis, H., Klee, K., Petrova, A., Wiegandt, R., Fust, A., Preussner, J., Kuenne, C., Braun, T., Kim, J. & Looso, M. ATAC-seq footprinting unravels kinetics of transcription factor binding during zygotic genome activation. Nature Communications 11, 4267, doi:10.1038/s41467-020-18035-1 (2020).

59 Heinz, S., Benner, C., Spann, N., Bertolino, E., Lin, Y. C., Laslo, P., Cheng, J. X., Murre, C., Singh, H. & Glass, C. K. Simple combinations of lineage-determining transcription factors prime cis-regulatory elements required for macrophage and B cell identities. Mol Cell 38, 576–589, doi:10.1016/j.molcel.2010.05.004 (2010).

60 Lindgreen, S. AdapterRemoval: easy cleaning of next-generation sequencing reads. BMC Research Notes 5, 337, doi:10.1186/1756-0500-5-337 (2012).

61 Patro, R., Duggal, G., Love, M. I., Irizarry, R. A. & Kingsford, C. Salmon provides fast and bias-aware quantification of transcript expression. Nature methods 14, 417 (2017).

62 Ritchie, M. E., Phipson, B., Wu, D., Hu, Y., Law, C. W., Shi, W. & Smyth, G. K. limma powers differential expression analyses for RNA-sequencing and microarray studies. Nucleic Acids Research 43, e47–e47, doi:10.1093/nar/gkv007 (2015).

63 Robinson, M. D., McCarthy, D. J. & Smyth, G. K. edgeR: a Bioconductor package for differential expression analysis of digital gene expression data. Bioinformatics (Oxford, England) 26, 139–140, doi:10.1093/bioinformatics/btp616 (2010).

64 Ramírez, F., Ryan, D. P., Grüning, B., Bhardwaj, V., Kilpert, F., Richter, A. S., Heyne, S., Dündar, F. & Manke, T. deepTools2: a next generation web server for deep-sequencing data analysis. Nucleic Acids Research 44, W160–W165, doi:10.1093/nar/gkw257 (2016).

65 Umansky, K. B., Gruenbaum-Cohen, Y., Tsoory, M., Feldmesser, E., Goldenberg, D., Brenner, O. & Groner, Y. Runx1 Transcription Factor Is Required for Myoblasts Proliferation during Muscle Regeneration. PLOS Genetics 11, e1005457, doi:10.1371/journal.pgen.1005457 (2015).

