## Supplementary Methods and Figures for "Progesterone receptor-A isoform interaction with RUNX transcription factors controls chromatin remodelling at promoters during ovulation"

**Quantitative real-time PCR**

21-day old female mice were hormonally stimulated by intraperitoneal injection with 5 IU eCG following by 5 IU hCG at 46 hours post-eCG. For time course gene expression, female CBA x C57BL/6 F1 (CBAF1) mice were culled at the following time points: 0 h (no hCG), 4 h, 6 h, 8 h, 10 h and 12 h post-hCG. Mice not treated with eCG and hCG were used as negative control (unstimulated). For PGRKO gene expression, female mice that were WT, het or KO from the PGRKO, AKO and BKO strains were culled at 8 h post-hCG. Granulosa cells from all mice were collected from dissected ovaries by repeated puncturing of the ovaries with a 26G needle in media. Granulosa cells from three mice were pooled together for RNA extraction. A total of three independent experimental replicates were conducted. RNA was extracted from granulosa cells using RNeasy Mini kit (Qiagen, Chadstone, VIC, Australia) for time course experiment or using Trizol RNA extraction method, including DNase treatment, for PGRKO experiment. cDNA was synthesised from 500 ng extracted RNA using SuperScriptIII Reverse Transcriptase kit (Thermo Fisher) and used for qPCR with Taqman methodology. Gene expression in each biological replicate (3 biological replicates per time point) was normalised to *Rpl19* and fold change was presented as relative to the mean of unstimulated samples using the ddC_T_ method. One way ANOVA was used as indicated in the figure legends and statistical significance was considered as p-value ≤ 0.05.

**Immunofluorescence imaging**

Superovulated CBAF1 female mice were culled and dissected at 0h (no hCG), 4 h, 6 h or 8 h post-hCG for intact ovaries. Tissues were fixed in 4% formaldehyde, embedded in paraffin and sectioned. Sections were dewaxed and rehydrated, then antigen retrieval was performed in citrate buffer (pH 6). Slides containing tissue sections were blocked for 1 h at room temperature in TBS with 10% goat serum, then incubated with primary antibodies (RUNX1 – ab23980, Abcam; RUNX2 – sc 390715, SCT) in 1:250-500 dilution and corresponding Alexa Fluor-conjugated secondary antibodies and Hoechst 33342 (nuclear staining) at 1:2000, each for 1 h at room temperature. Slides were then mounted with mounting media (Dako, Santa Clara, USA) and imaged using the Olympus FV3000 confocal laser scanning microscope (Olympus, Tokyo, Japan).

**Ovulation rate assessment**

Female mice that are WT, het and KO from the PGRKO, AKO and BKO strains were hormonally stimulated as above. Mice were humanely killed by cervical dislocation 16 h post-hCG and the oviducts were dissected to obtain ovulated cumulus-oocyte complexes (COCs). Results are presented as the number of ovulated COCs per animal. One-way analysis of variance (ANOVA) was used as indicated in the figure legends and statistical significance was considered as p-value ≤ 0.05.

**Supplementary table 1: TOBIAS-identified differential binding motifs**

**Supplementary table 2: RNA-seq DEG (LH: 8h-hCG vs 0h-hCG, PGRKO: WT vs KO, AKO: WT vs KO)**

**Supplementary table 3: Upstream regulators of LH RNA-seq**

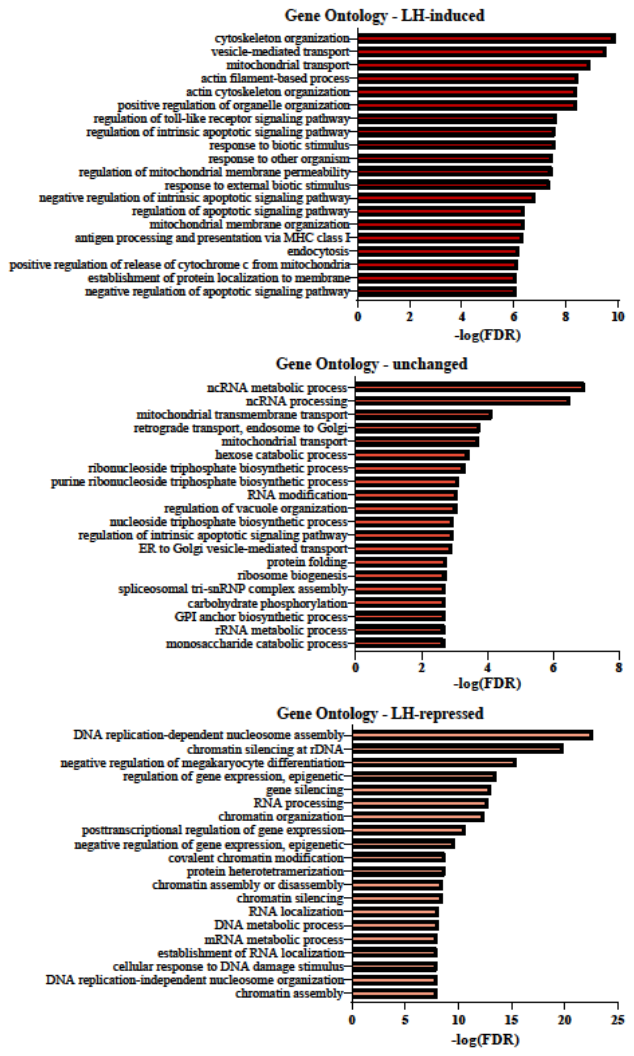
**Supplementary figure 1:** GREAT analysis for GO Biological Process enrichment of ATAC-seq peaks that were LH-induced (top), unchanged (middle) and LH-repressed (bottom). For each subset, only peaks in promoter region (within 3 kb upstream of TSS) were analysed and the top 20 terms are listed.

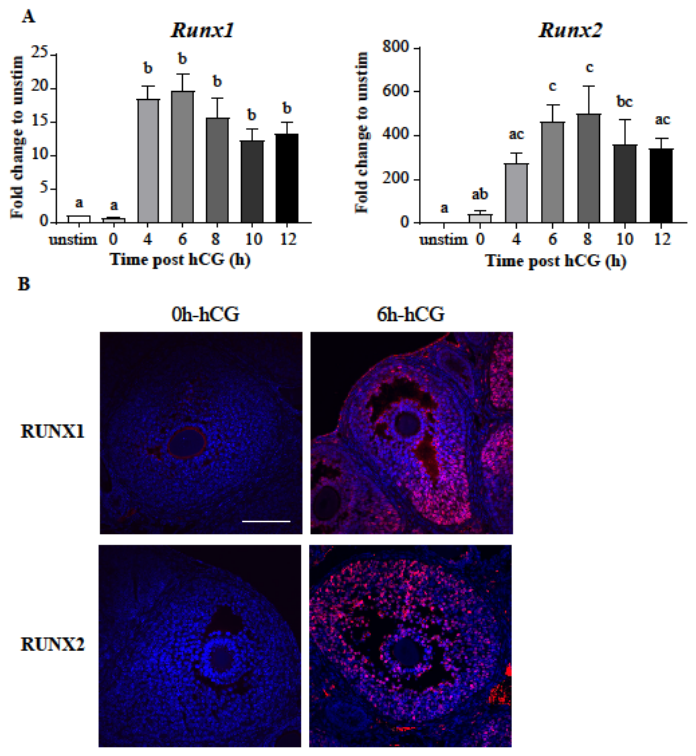

**Supplementary figure 2:** (A) Expression of *Runx1* and *Runx2* mRNA in peri-ovulatory granulosa cells as determined through RT-qPCR. RT-qPCR was performed on granulosa cells obtained either without eCG or hCG (unstim) or eCG + hCG 0-12 h stimulation. Fold change is displayed as normalised to the reference gene *Rpl19* and relative to the unstim sample, mean ± SEM, N = 3 biological replicates (3-5 mice per replicate per time point). Bars with different superscripts are statistically different (p-value < 0.05). (B) RUNX1 and RUNX2 immunofluorescence in ovarian sections obtained from eCG-mice stimulated with hCG for 0 h or 6 h. Antibody labelling is in red and nuclear staining in blue. Scale bar = 100 µm.

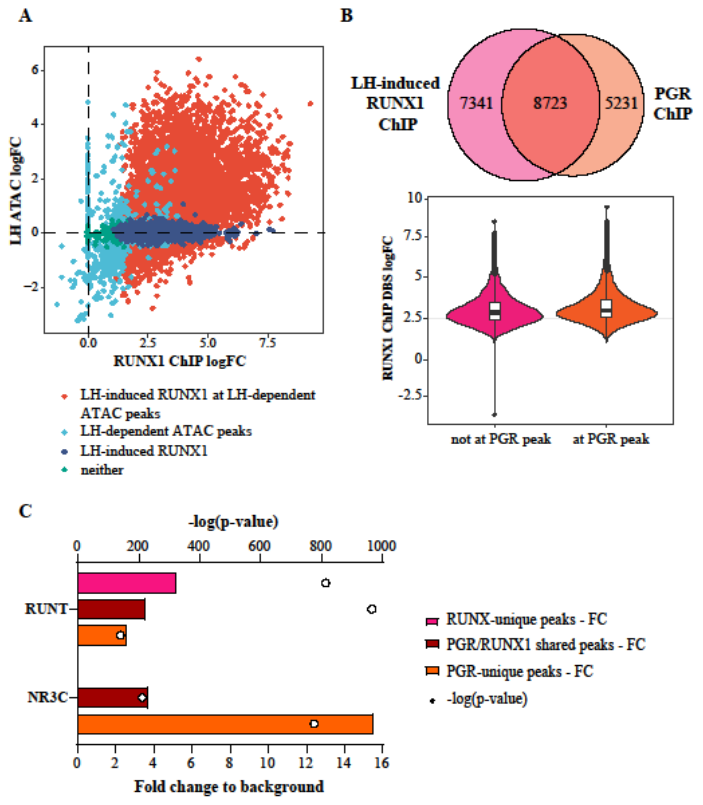
**Supplementary figure 3:** (A) Scatter plot of LH-induced RUNX1 ChIP-seq fold change vs ATAC fold change at same site. Colour label corresponds to sites identified as differentially enriched RUNX1 sites, LH-dependent ATAC sites or both. (B) Level of LH-induced RUNX1 binding at chromatin sites with or without PGR-cobinding, with Venn diagram showing the number of shared sites and violin plot showing the distribution of RUNX1 ChIP-seq fold change. (C) Enrichment level of the RUNT and NR3C motifs at binding sites specific to RUNX1 (pink), PGR (orange) or shared (brown). Bars indicate fold enrichment of motif to background (bottom x axis). Circles indicate -log(p-value) (top x axis).

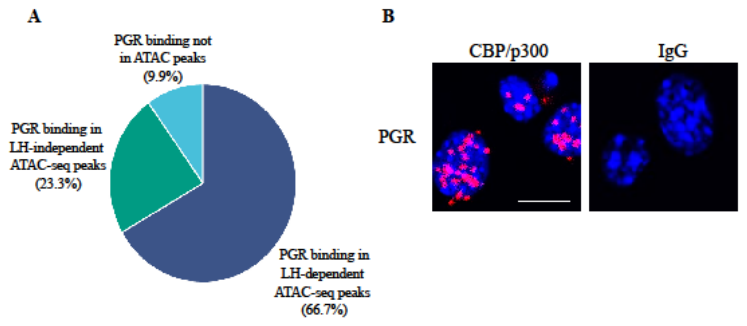
**Supplementary figure 4:** (A) The proportion of PGR binding sites in LH ATAC-seq peaks, divided into PGR binding sites in LH-dependent ATAC sites, in LH-independent ATAC sites or not in ATAC-defined sites. (B) PLA showing the interaction between PGR and acetyl-CBP/p300 or IgG (negative control) in granulosa cells treated for 6 h with hCG and R5020. Scale bar = 10 µm.

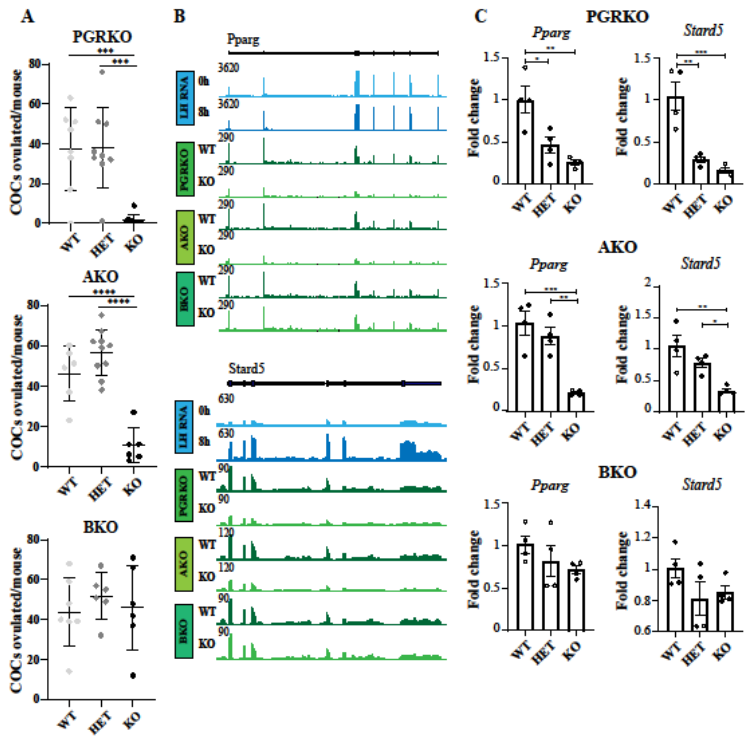
**Supplementary figure 5:** (A) Ovulation rate of female mice that are WT, het or KO for total PGR (PGRKO), PGR-A (AKO) or PGR-B (BKO). (B) Examples of LH RNA-seq (track 1, 2), PGRKO RNA-seq (track 3, 4), AKO RNA-seq (track 5, 6) and BKO RNA-seq (track 7, 8) signals at *Pparg* and *Stard5*. (C) qPCR of *Pparg* (left) and *Stard5* (right) in granulosa from WT, het and KO mice in PGRKO (top), AKO (middle) and BKO (bottom) strains. Fold change is displayed as normalised to the reference gene *Rpl19* and relative to WT, mean ± SEM, * = p-value < 0.05, ** = p-value < 0.01, *** = p-value < 0.0005, **** = p-value < 0.0001. N = 4 biological replicates (3-5 mice per replicate per genotype).
